# Regulation of Exocytosis and Mitochondrial Relocalization by Alpha-Synuclein in a Mammalian Cell Model

**DOI:** 10.1101/492066

**Authors:** Meraj Ramezani, Marcus M. Wilkes, Tapojyoti Das, David Holowka, David Eliezer, Barbara Baird

## Abstract

We characterized phenotypes in RBL-2H3 mast cells transfected with human alpha synuclein (a-syn) using stimulated exocytosis of recycling endosomes as a proxy for similar activities of synaptic vesicles in neurons. We found that low expression of a-syn inhibits stimulated exocytosis and that higher expression causes slight enhancement. NMR measurements of membrane interactions correlate with these functional effects: they are eliminated differentially by mutants that perturb helical structure in the helix 1 (A30P) or NAC/helix-2 (V70P) regions of membrane-bound a-syn, but not by other PD-associated mutants or C-terminal truncation. We further found that a-syn (but not A30P or V70P mutants) associates weakly with mitochondria, but this association increases markedly under conditions of cellular stress. These results highlight the importance of specific structural features of a-syn in regulating vesicle release, and point to a potential role for a-syn in perturbing mitochondrial function under pathological conditions.

## INTRODUCTION

Parkinson’s disease (PD) is the second most common neurodegenerative disorder.^1^ With increased risk of diagnosis after age 60, PD onset strongly correlates with age, affecting ~1-2% of the population over age 65.^2^ Clinically, motor dysfunction in PD is associated with death of dopaminergic neurons in the substantia nigra, coupled with the presence of abnormal protein aggregates, known as Lewy bodies, in surviving neurons.^2,3^ The primary component of these intraneuronal aggregates is the presynaptic protein alpha synuclein (a-syn).^4^ A-syn inclusions are also a defining feature of Parkinson’s Disease Dementia and Lewy Body Dementia, as well as a common co-pathology in Alzheimer’s disease.^5^

First identified as binding to synaptic vesicles,^6^ a-syn has been characterized as a 140 amino acid, intrinsically disordered protein found predominantly in neurons.^7^ A-syn is genetically linked to autosomal dominant early onset familial PD through point mutations and gene duplication or triplication.^8–10^ Elevated levels of a-syn are observed in many patients who develop sporadic forms of PD, suggesting a critical role for this protein in the majority of PD cases.^11,12^ Moreover, genome wide-association studies have identified the gene that encodes for a-syn, SNCA, as one of the strongest risk loci for sporadic forms of PD.^13^ A-syn has been the subject of intense research with the aim of understanding the relationship between a-syn physiological function and PD pathology, but this connection remains poorly understood at the cellular level.

Several studies with a-syn in neuronal cells and synaptosomes derived from rodent brains implicate a critical role in regulation of SV trafficking (exocytosis and endocytosis) and homeostasis,^14^ and membrane binding properties related to a-syn structure have been evaluated *in vitro*.^15,16^ The amphipathic N-terminal segment (residues 1-100; Figure 1A) forms an extended helix upon binding to negatively charged phospholipid vesicles (Figure 1B), which breaks into two smaller helices when binding to phospholipid micelles (Figure 1C). The broken helix form has been proposed to resemble the structure of a-syn when binding simultaneously to two phospholipid membranes, such as bridging a synaptic vesicle with the plasma membrane.^17^ The acidic C-terminal segment of a-syn (residues 100-140) is disordered and suppresses a-syn aggregation mediated by the N-terminal NAC region (Figure 1A).

**Figure 1.**
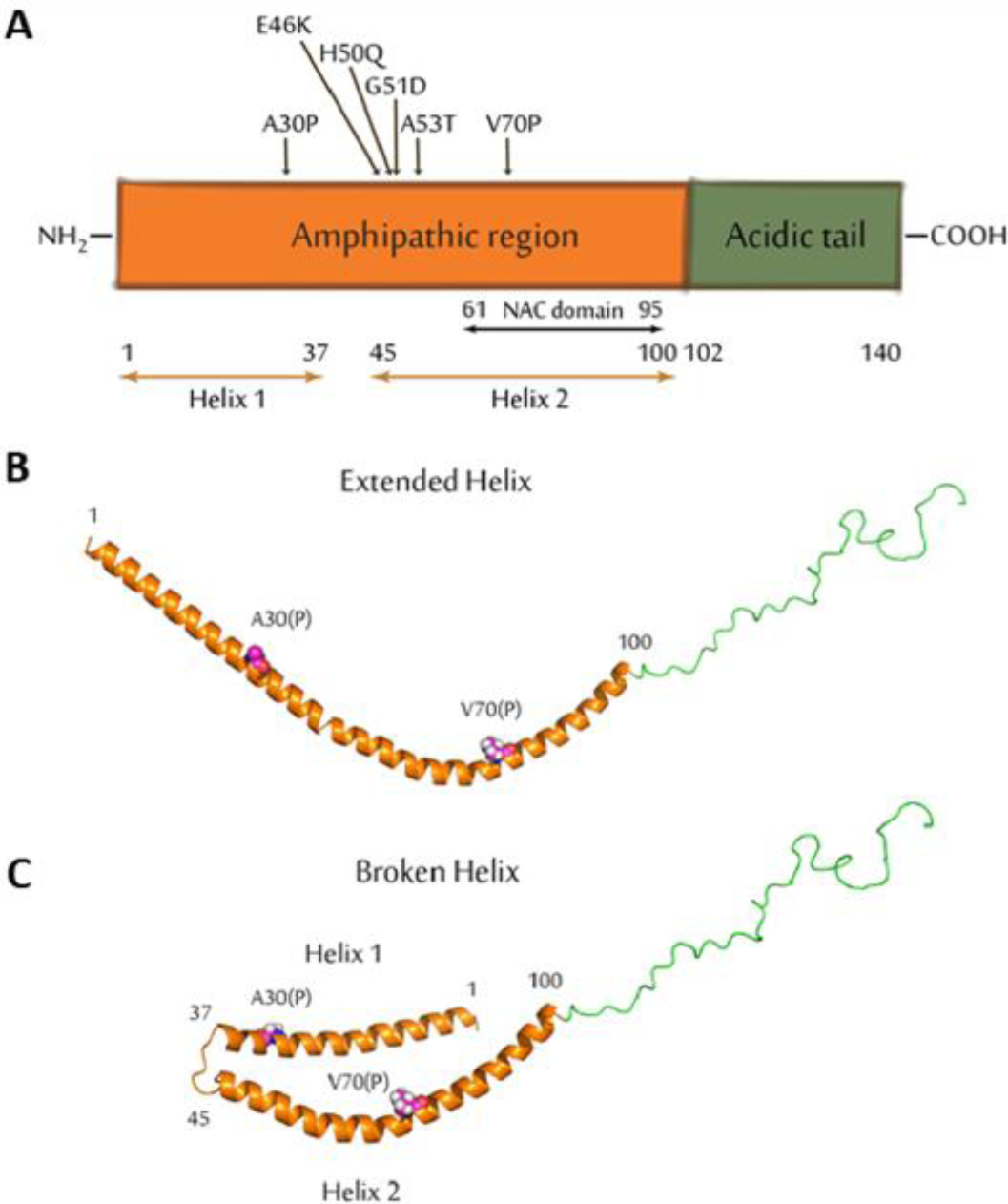
Structural features of a-syn. **A**) Schematic representation of the a-syn primary sequence delineating the amphipathic membrane-binding domain (orange) and the acidic Cterminal tail (green), and indicating the locations of the NAC domain, helix-1 and helix-2 of the broken-helix state, and the sites of PD and other mutations examined in the manuscript. **B**) Cartoon model of Wt a-syn in an extended helix conformation. The model was generated from RCSB protein data bank entry 1XQ8 by manually converting the non-helical linker between helix-1 and helix-2 to a helical conformation. The sidechains of residues Ala 30 and Val 70 are shown to indicate the location of proline mutations examined in this study. **C**: Wt a-syn in the broken helix conformation (RCSB protein data bank entry 1XQ8) with the sidechains of Ala 30 and Val 70 shown.

Delineating the structural mechanisms by which a-syn regulates SV trafficking, and if and how this becomes dysregulated, is necessary for understanding PD onset and pathology. Toward this objective, we developed an experimental system based on trafficking and stimulated exocytosis of recycling endosomes (REs) in RBL-2H3 mast cells, which have proven useful as a model system for investigating general questions of membrane and cell biology. Our preliminary studies demonstrated a clear phenotype: Human a-syn expressed at low levels in these cells inhibits stimulated exocytosis of REs. With this result as a starting point we systematically compared Wt a-syn to a panel of a-syn variants, including a-syn mutants associated with PD and a-syn with mutations in N-terminal helical and C-terminal tail regions (Figure 1). We correlated functional effects with effects on binding modes to model membranes *in vitro*, quantified using NMR measurements. Surprisingly, we found that whereas low expression levels of Wt a-syn inhibit stimulated RE exocytosis, high expression levels have an enhancing effect. Moreover the effects of specific a-syn mutants are different for high vs low expression levels. These differential effects prompted us to evaluate intracellular distributions of a-syn and REs using fluorescence imaging, and we observed that high level expression of a-syn causes dispersal of REs from the endocytic recycling compartment (ERC), which may play a role similar to the reserve pool of SVs in neurons. We also observed a-syn association with mitochondria, depending on expression level, mutations, and mitochondrial stress. Mitochondrial dysfunction and stress are strongly implicated in PD by genetics (e.g. PINK/parkin pathway) and environmental factors (e.g. pesticides), and stress-dependent association of a-syn with mitochondria could provide a link between these two different critical pathways in PD.

Overall, our results provide a self-consistent picture of how a-syn affects the functions of RBL cells that can be related directly to modes of a-syn binding to synthetic membranes as measured with NMR. The picture emerging for a-syn effects is that stimulated RE exocytosis is: inhibited by high affinity binding to docked vesicles via a broken helix, requiring an intact helix 2 (Figure 1C); and enhanced at high expression levels by curvature sensitive, low affinity binding to isolated vesicles/tubules involving helix 1 (Figure 1B), leading to increased availability of vesicles for docking. Specific membrane binding properties also provide insight to variable association of Wt a-syn and mutants with mitochondria with and without stress. We point to the parallels between trafficking of endosomal vesicles and association with internal organelles in neurons and non-neuronal cells, and suggest that the a-syn interactions and effects we have characterized with REs in mast cells may be related to the interactions of asyn with SVs and mitochondria within neurons that are implicated in the etiology of PD.

## Results

### RBL-2H3 mast cells serve as a model for stimulated exocytosis and related signaling events

Because of the difficulties of examining a-syn participation in neuronal function and dysfunction in primary neurons and because there is currently a paucity of related cell biological information, we developed the RBL model cell system for investigating general structure-function relationships using a panel of a-syn mutants. RBL cell responses are typically triggered via antigen crosslinking of immunoglobulin E bound to its high affinity receptor (IgE/FcεRI), which results in exocytosis of both secretory lysosomes (degranulation) and recycling endosomes (REs).^18,19^ However, just as neuronal release of SVs involves increase of cytosolic Ca^2+^ concentration,^20^ so does stimulated exocytosis in RBL cells involve Ca^2+^ mobilization that can be triggered directly with thapsigargin, which inhibits the sarco/endoplasmic Ca^2+^ ATPase.^21^ Also consistent with this approach, a previous study suggested that a-syn inhibition of dopamine exocytosis in PC12 and chromaffin cells occurs downstream of stimulated Ca^2+^ responses.^22^

### Wt a-syn interferes with stimulated exocytosis in RBL cells

We showed previously that stimulated exocytosis of REs is triggered by both antigen and thapsigargin.^23^ Total internal reflection microscopy (TIRFM) shows vividly that low expression levels of Wt a-syn inhibit thapsigargin-stimulated exocytosis (Figure 2A,B and supplemental Movies S1A,B). To view this time course by microscopy, cells were co-transfected with DNA for human a-syn or vector control (pcDNA), together with the reporter VAMP8-pHluorin, a v-SNARE protein conjugated to a fluorophore that is quenched when localized to the mildly acidic environment of the RE. VAMP8-pHluorin fluorescence increases markedly after stimulated trafficking of REs to, and exocytosis at, the plasma membrane, thereby encountering the neutral pH environment of the extracellular medium.^23^ Addition of NH^4+^Cl^−^ at the end of the time course neutralizes the pH in all cellular compartments and provides a means of normalizing the level of the stimulated exocytosis. TIRFM further reveals that, in contrast to lower a-syn expression levels which inhibit exocytosis, cells expressing higher levels of a-syn exhibit a faster rate and overall greater extent of exocytosis compared to pcDNA controls (Figure 2C, Movie S1C). The following sections quantify these opposing effects of a-syn at lower and higher expression levels, and selected a-syn mutants are evaluated for additional insight into the structural interactions involved. Table 1 summarizes key results.

**Figure 2.**
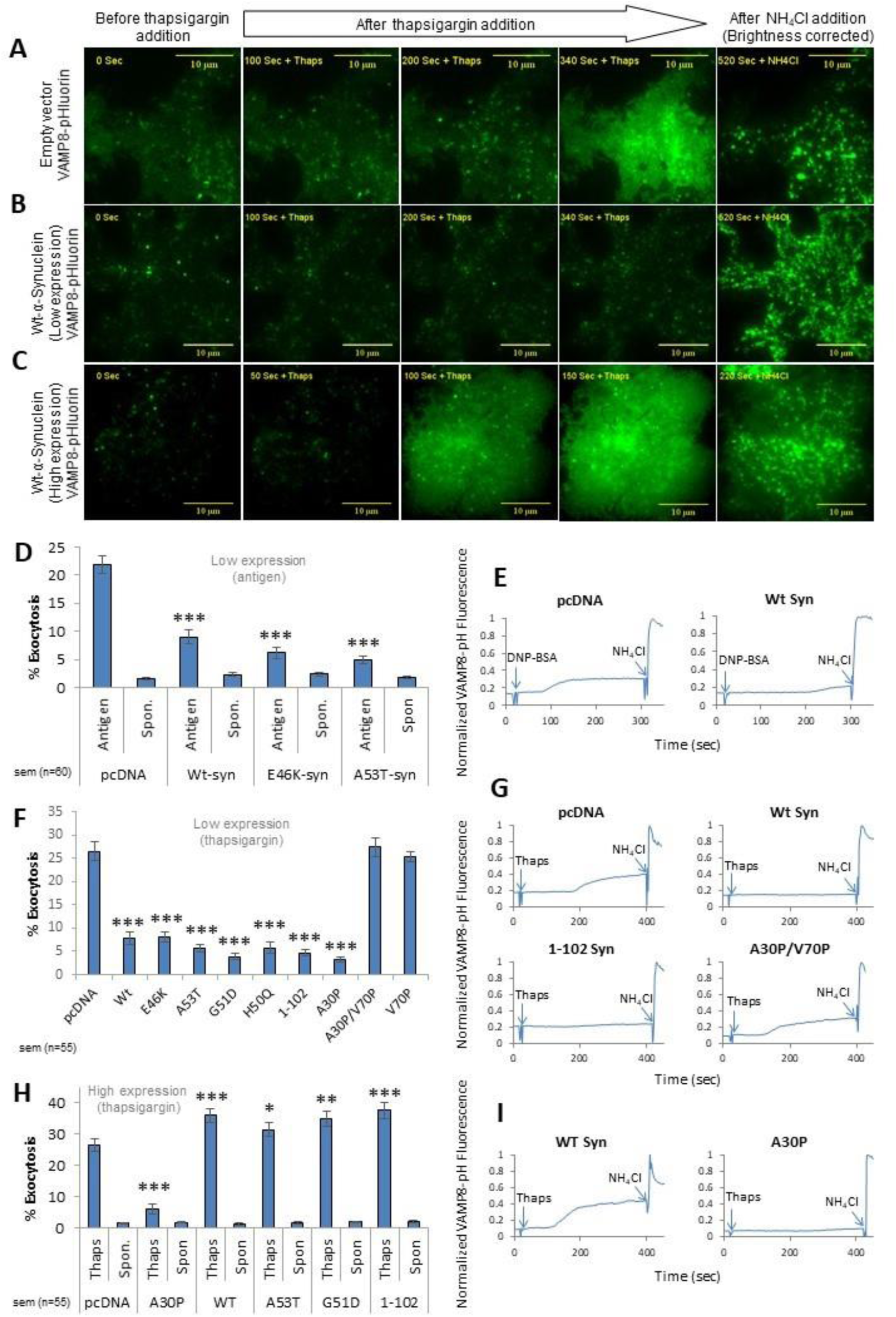
Low expression levels of Wt a-syn inhibit stimulated exocytosis of REs, whereas high expression levels cause enhancement, and mutants of a-syn show distinct differences. As indicated, RBL cells were co-transfected with VAMP8-pHluorin and low or high levels of pcDNA, Wt a-syn or selected mutants of a-syn. Exocytosis was stimulated by thapsigargin (250 nM; **A**-**C**, **F**-**I**) or DNP-BSA (1 ng/ml; **D**, **E**). VAMP8-pHluorin fluorescence increase was monitored in movies (Supplemental Movies SM 1-5) before and after stimulation, and after indicated addition of NH_4_Cl (50mM, 300-400 sec later) to dequench intracellular VAMP8-pHluorin fluorescence. **A**-**C)** Snapshots from representative movies taken with TIRFM (Movies SM1-3); scale bar = 10μm. Brightness of images in right-most column was reduced by 50% for better detail visualization. **D)** Averaged exocytosis stimulated by DNP-BSA in cells with low a-syn expression levels (n=60). **E)** Representative traces of VAMP8-pHluorin fluorescence integrated from movies of multiple confocal fields of 5-6 cells (all traces shown in Figure S8, Movies SM 4-5). **F)** Averaged exocytosis stimulated by thapsigargin in cells with low a-syn expression levels (n=55), spontaneous release for each of these samples is less than 2.5% as shown in Figure S9. **G)** Representative traces of VAMP8-pHluorin fluorescence for **F** as in **E** (all traces shown in Figure S9). **H)** Averaged exocytosis stimulated by thapsigargin in cells with high a-syn expression levels (n=55). **I)** Representative traces of VAMP8-pHluorin fluorescence integrated from movies as in **E** (All traces shown in Figure S10) All data sets shown are from 34 independent experiments; Error bars are ± SEM; ^***^ represents P-values <0.001, ^**^ represents P-values <0.01, ^*^ represents P-values <0.05.

**Table 1.** Functional effects and structural interactions of Wt a-syn and key mutants. ↑ and ↓ indicate enhancing or inhibitory effect, respectively; +++, High effect; +, Low effect; --, No effect; NT, not tested.

| Conditions | Wt-a-syn | A30P-a-syn | V70P-a-syn |
| --- | --- | --- | --- |
| Stimulated Exocytosis - Low expression (Fig. 2F) | ↓+++ | ↓+++ | -- |
| Stimulated Exocytosis - High expression (Fig. 2H) | ↑+ | ↓+++ | ↑+ |
| Binding to Mitochondria - No stress (Fig. 6) | + | -- | -- |
| Binding to Mitochondria - With stress (Fig. 6) | +++ | + | -- |
| Binding to Lipid droplets - High expression levels (Fig. S6) | +++ | -- | NT |
| Disruption of vesicle binding(SUVs, NMR) (Fig. 3,S3) | Helix1— Helix2<br>-- -- | Helix1— Helix2<br>+ +++ | Helix1— Helix2<br>-- +++ |
| Disruption of secondary structure (micelles, NMR) (Fig. 3) | Helix1— Helix2<br>-- -- | Helix1— Helix2<br>-- + | Helix1— Helix2<br>-- + |

### Wt, A53T, and E46K a-syn expressed at low levels inhibit antigen-stimulated exocytosis of REs

We quantified the normalized change in VAMP8-pHluorin fluorescence using confocal microscopy. RBL cells co-transfected with DNA for Wt a-syn or PD-linked a-syn mutants A53T and E46K and sensitized with anti-DNP IgE were stimulated with DNP^16^BSA. The DNP-antigen crosslinks IgE/FceRI on the cell surface to initiate a signaling cascade, activating protein kinase C (PKC) and triggering Ca^2+^ release from intracellular stores, followed by store operated Ca^2+^ entry from the extracellular medium and consequent exocytosis of secretory lysosomes and REs.^23^ Cells transfected with DNA for pcDNA, together with VAMP8-pHluorin, and stimulated with a low dose of DNP^16^BSA (1 ng/ml) exhibit increased fluorescence at the plasma membrane (Figures 2E, S8A; Movie S2A). However, low levels of Wt a-syn coexpressed with VAMP8-pHluorin strongly inhibit this stimulated exocytosis (Figures 2E, S8B; Movie S2B) as do a-syn mutants A53T and E46K (Figures S8E, F). Results quantified for multiple experiments show averaged inhibition ranging from 70% to 85% for these three constructs, compared to pcDNA controls (Figure 2D).

We previously determined that VAMP8-phluorin is selective for REs.^23^ VAMP7-pHluorin serves as a secretory granule/lysosome marker,^24,25^ and Wt a-syn may slightly inhibit antigen stimulated degranulation (Figure S8H). However, we don’t find this decrease to be statistically significant, indicating that the significant inhibition by a-syn we observe under these conditions is selective for stimulated exocytosis of REs. We also tested whether inhibition of exocytosis by a-syn is detectable with higher-dose antigen stimulation and determined that it is not: Stimulating co-transfected RBL cells first with a low dose of antigen (1 ng/ml DNP-BSA) shows inhibition, but this is abrogated if followed by a higher dose (200 ng/ml DNP-BSA) as compared to cells with the vector control (Figure S8I,J). These results show that higher antigen doses overcome the inhibitory mechanism and also that the a-syn inhibition observed with low dose antigen is not due to cytotoxicity.

### Low level expression of Wt a-syn, PD-linked and C-terminal mutants, but not V70P mutants, inhibit thapsigargin-stimulated exocytosis of REs

We quantified inhibitory effects of Wt and a broad range of a-syn mutants with confocal microscopy, using thapsigargin to stimulate Ca^2+^ mobilization. As shown in Figure 2G, thapsigargin stimulates RE exocytosis in RBL cells, as reported by VAMP8 pHluorin in vector control samples (consistent with Figure 2A). At low expression levels, Wt, A53T, and E46K a-syn all inhibit thapsigargin-stimulated exocytosis of REs (Figures 2F and S9B-D), by 70-85%, similar to that observed for antigen stimulation (Figures 2D and S8D-F), as do PD-linked a-syn mutants, H50Q and G51D (Figure 2F).

Previous studies reported that the disordered C-terminus of a-syn (Figure 1) mediates interactions with proteins involved in exocytosis, including the v-SNARE VAMP2^26^ and Rab family proteins,^27^ which also participate in stimulated exocytosis of REs in RBL cells.^23^ We found that truncated a-syn 1-102 (C-terminal residues 103-140 deleted) inhibits thapsigargin mediated exocytosis similarly to Wt a-syn (Figure 2F,G). We also tested Wt a-syn tagged with mRFP on its C-terminus (Wt syn-mRFP), considering that a fluorescent tag the size of mRFP (~27 kDa) may sterically disrupt some cellular interactions of a-syn (~14 kDa) and found it to be similarly inhibitory (Figure S1B, E). These results indicate that the C-terminus is not involved in the inhibition of stimulated exocytosis we observe at low expression levels of Wt a-syn.

We evaluated the contributions of selected residues within the N-terminal amphipathic region of a-syn (Figure 1), which has been shown to be necessary for membrane binding.^28^ The PD-linked mutant A30P perturbs the first of two alpha helical regions formed in this region^29,30^ and decreases membrane affinity.^31^ A30P a-syn at low expression levels inhibits stimulated exocytosis under our conditions (Figure 2F and S9E). To further reduce a-syn membrane binding, we introduced a valine to proline mutation at residue 70 within the A30P mutant (A30P/V70P). The V70P mutation is located in the second helical region of membrane bound a-syn (Figure 1) and was previously shown to decrease membrane binding when combined with an alanine to proline mutant at position 11 (A11P/V70P).^28,32^ Interestingly, A30P/V70P a-syn mutant does not inhibit thapsigargin-stimulated exocytosis under the same conditions (Figures 2F,G), indicating that some aspect of membrane binding is essential for the inhibition we observe. We then examined the single V70P mutation and observed that it too fails to inhibit thapsigargin-stimulated exocytosis (Figures 2F and S9G), indicating that perturbations in the helix-2 region of membrane-bound a-syn are sufficient, and A30P is not required, to disrupt its inhibitory effect.

### The V70P mutation perturbs membrane binding affinity of a-syn helix-2, but not of helix1

To assess the structural consequences of the A30P and V70P mutations, alone or in combination, for membrane-bound a-syn we carried out NMR measurements on the micelle bound and lipid vesicle-bound forms of the protein. The vesicle-bound state of a-syn is expected to closely resemble the state of the protein when bound to undocked synaptic vesicles *in vivo*.^16,33,34^ The micelle-bound state of the protein has no direct physiological analogue, but we and others have postulated that this state resembles the structure of a-syn when it is bound simultaneously to two different membrane surfaces, for example in a mode bridging the vesicle and plasma membranes, or potentially bridging two adjacent synaptic vesicles.^16,17,35^ Analysis of ^1^H/^15^N chemical shift changes compared to Wt a-syn indicates that the V70P mutation alters the environment of sites as far away as 15 residues from the site of the mutation in the micelle-bound state (green trace, Figure 3A). Deviations of Ca shifts from random coil values, which are indicative of secondary structure (green trace, Figure 3B), indicate that helix-1 is unperturbed in the V70P mutant, whereas helix-2 exhibits reduced helicity around the site of the mutation, with a severe reduction of helical structure spanning about 10 residues centered on the site of the mutation. Together, these changes indicate that the V70P mutation does not perturb membrane binding associated with helix-1, but does locally perturb and deform the membrane-bound conformation of helix-2.

**Figure 3.**
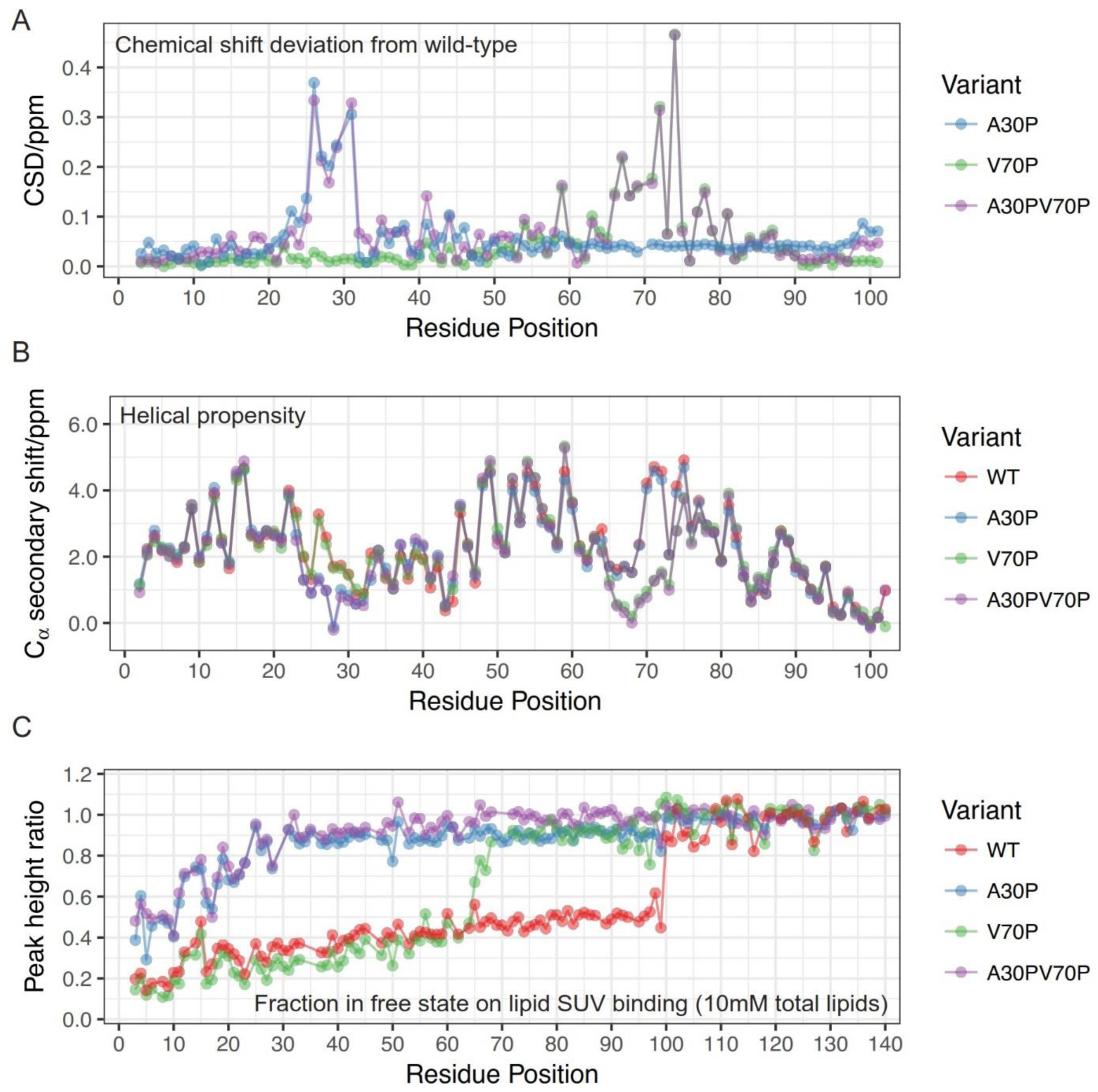
The V70P mutation perturbs helical micelle-bound structure locally and results in release of a-syn regions C-terminal to the mutation site from vesicle membranes. **A**) Average amide chemical shift deviation (CSD) of different a-syn mutants used in the study from Wt a-syn, all in the micelle-bound state, calculated as described in Methods. Only the N-terminal 102 residues are plotted as the C-terminal region does not interact with the micelle. **B**) C_α_ secondary chemical shift of a-syn variants in the micelle-bound state, calculated for each residue as the difference between the observed chemical shift and the tabulated chemical shift in a random coil conformation. Positive values indicate helical structure. **C**) Vesicle binding of full-length a-syn variants measured as the ratio of NMR resonance intensities in the presence and absence of liposomes. The peak intensity ratio, representing the free fraction of each residue, is plotted for 50μM protein with small unilamellar vesicles (SUVs) containing 10mM total phospholipids at a molar ratio of DOPC:DOPE:DOPS = 60:25:15.

The vesicle-bound state of a-syn cannot be directly visualized using solution state NMR because of the slow tumbling time of lipid vesicles. Instead, NMR allows for observation of any parts of the protein that are not bound to the vesicle surface (and are therefore highly mobile). For any given signal originating from a specific residue site in the protein, the intensity of the signal, normalized by the intensity of the same signal in the absence of lipids, indicates the fraction of the protein population for which that protein residue is not bound to the vesicle surface. For Wt a-syn, these intensity ratio plots demonstrate that the entire membrane-binding domain of the protein, consisting of residues ~1-100, bind to the vesicle surface largely as a single unit (red traces, Figures 3C and S3). Indeed, more detailed structural studies have indicated that in this binding mode, helix-1 and helix-2 fuse into a single extended helix (Figure 1B).^36–39^ For the V70P mutant, binding to the membrane surface is dramatically reduced for locations C-terminal to the site of the mutation, whereas binding for locations N-terminal to the mutation site is largely unaffected, and, in fact, appear to be slightly enhanced (green traces, Figures 3C and S3). This indicates that the proline at position 70 leads to detachment from the membrane surface of the portion of the protein following this mutation site, but not of the preceding portion.

We compared the effects on micelle and vesicle binding of the V70P mutation to those of the A30P mutation (blue traces) and of the A30P/V70P double mutant (purple traces) (Figures 3 and S3). As previously reported,^29,30^ the A30P mutation locally perturbs the structure of helix-1 in the micelle-bound state (blue trace, Figure 3A), and leads to decreased vesicle binding of a-syn residues both N-terminal to and C-terminal to the site of mutation (blue trace, Figure 3C), consistent with the reduced membrane affinity of this mutant. The effects of the double A30P/V70P mutation appear to be decoupled and additive in the micelle-bound state, with both helix-1 and helix-2 exhibiting local perturbation of helical structure, but with the micelle-bound structure remaining otherwise intact (purple trace, Figure 3A, B). In contrast, in the vesicle-binding assay, the effects appear to be cumulative, with a dramatic loss of binding around the A30P site, and an additional small decrease in binding occurring C-terminal to the V70P site (purple trace, Figures 3C and S3). Our collective results show that the effects of the A30P mutation dominate the binding of a-syn to isolated vesicles, yet this mutant inhibits stimulated exocytosis similarly to Wt, while V70P does not. Thus, it appears that the disruption of helix-2 structure by V70P in the micelle-bound state, which may reduce its capacity to bridge between different membranes or possibly perturb another functional interaction, underlies the loss of its ability to inhibit stimulated exocytosis.

### High expression of Wt a-syn enhances stimulated exocytosis of REs

To evaluate the dependence of a-syn function on expression levels we transfected RBL cells with the same amount of DNA for VAMP8-pHluorin as in Figure 2G, together with five times the amount of pcDNA or DNA for a-syn constructs. Cells expressing high levels of Wt, A53T, G51D, and 1102 human a-syn show robust thapsigargin-stimulated exocytosis (Figures 2H, I and S10C-G). Further, these responses show consistent enhancement over the thapsigargin-stimulated exocytic response observed with transfection of low levels of pcDNA alone (Figure 2H). Exocytosis data for cells transfected at low expression levels (Figure 2F) and high expression levels (Figure 2H) of a-syn constructs were typically collected on the same day. We used pcDNA transfected at low levels as a positive control, consistently exhibiting stimulated exocytosis values of about 25%. Expression at high levels of Wt, A53T, G51D, and 1-102 asyn increase this value by 20%-40% (Figure 2H). pcDNA transfected at higher concentrations gave somewhat variable results but consistently showed stimulated exocytosis less than that with the lower concentration of pcDNA. Wt a-syn-mRFP also loses the capacity to inhibit exocytosis when expressed at higher concentrations and shows a 20% increase in stimulated exocytosis when compared to mRFP alone expressed at high levels (Figure S1C-E). These results point to expression level-dependent interactions of a-syn: For multiple constructs, low expression levels cause inhibition, whereas high expression levels cause enhancement, of thapsigargin-stimulated exocytosis of REs.

In contrast to Wt and other mutants tested at high expression levels, A30P a-syn does not exhibit loss of inhibition as compared in the same experiment (Figures 2H, I and S10). In a separate experiment we found that V70P, which does not inhibit exocytosis at low expression levels (Figures 2F), also slightly enhances exocytosis at high expression levels (data not shown). Thus, it appears that while the interaction mode of a-syn with isolated vesicles is not critical for the capacity of a-syn to inhibit exocytosis, this interaction mode, and/or the structure of helix-1, both of which are perturbed by the A30P but not by the V70P mutation, may be important for stimulating vesicle exocytosis at higher expression levels.

### High expression levels of Wt a-syn cause dispersal of REs from endocytic recycling compartment (ERC)

To investigate possible explanations for differential effects on stimulated exocytosis at different a-syn expression levels we transfected cells with Wt a-syn or EGFP (as a control) together with mCh-Rab11, which labels REs.^23^ As shown in Figure 4 we observed markedly differential effects on the distribution of REs: Higher, but not lower, expression levels of Wt a-syn cause redistribution of the REs away from the perinuclear region of the cell, corresponding to the ERC, and toward the plasma membrane. Quantifying this redistribution as the ratio of membrane proximal to total Rab11 fluorescence, the value for Wt a-syn at high expression levels is 41% compared to 26% for Wt a-syn at low expression levels or 25% for EGFP at high expression levels, both highly significant differences (Figure 4B). Moreover, the value for A30P a-syn at high expression levels is similar to that for Wt a-syn at low expression levels. These differential effects on RE distribution are thus parallel to those observed for stimulated RE exocytosis, particularly our observation that high levels of Wt but not A30P asyn cause enhancement. It appears that proximity of abundant REs to the plasma membrane may underlie the enhancing effect.

**Figure 4.**
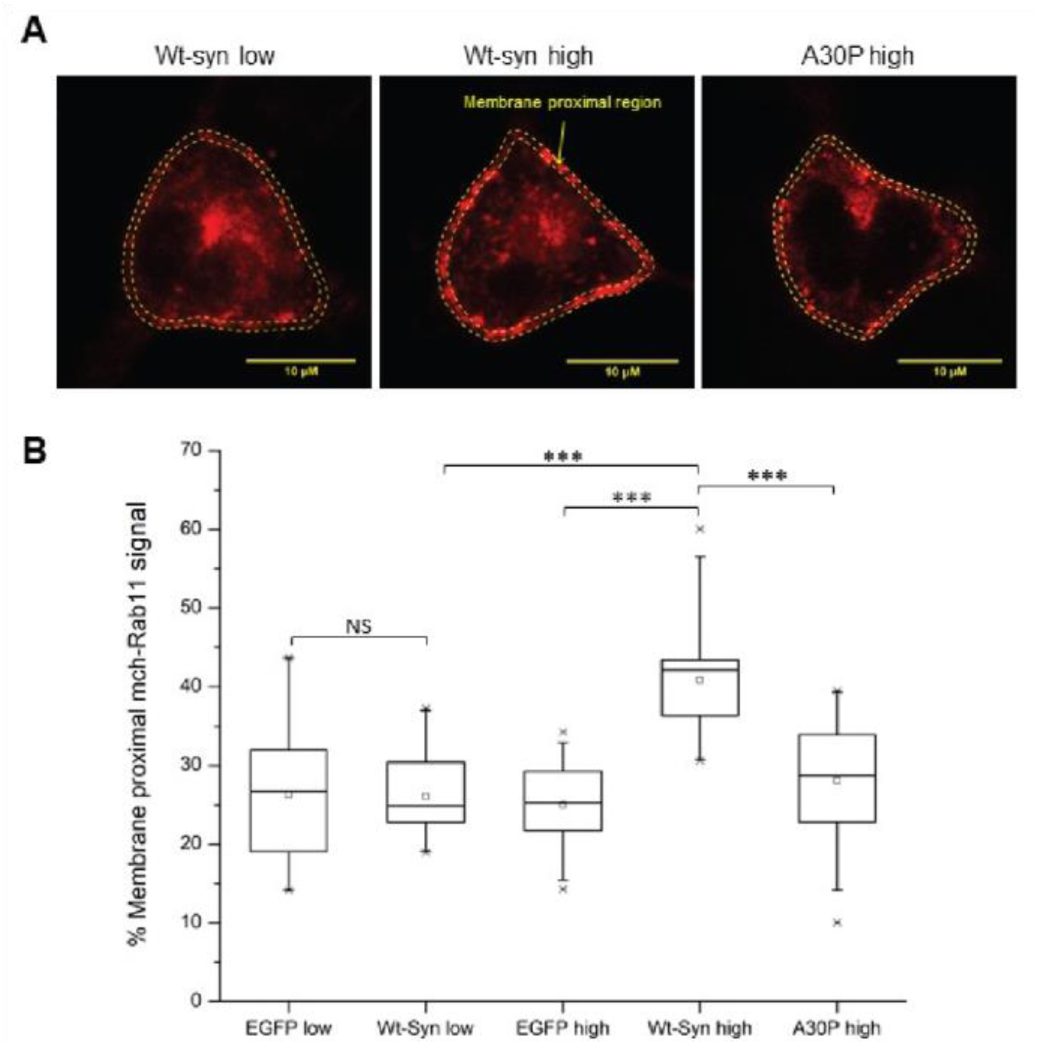
High expression levels of Wt, but not A30P, a-syn shift the distribution of REs toward the plasma membrane. RBL cells were co-transfected with mCh-Rab11 to label REs, and low or high levels of EGFP or Wt a-syn, or high levels of A30P a-syn. **A)** Confocal micrographs of mCh-Rab11 fluorescence (REs) co-transfected with low levels of Wt a-syn or high levels of Wt or A30P a-syn. The fluorescence intensity within a thin layer around the plasma membrane (dashed lines) divided by total cell fluorescence was used to calculate % membrane proximal REs; scale bar = 10μm. **B)** Averaged % membrane proximal REs for specified samples (about 20 cells for each condition) in a box plot. The box represents 25th75th percentile of the data, the midline represents the median and the small square represents the average. ^***^ represents P-values <0.001; NS, not statistically significant (P-values >0.05).

### Low and high expression levels of human a-syn are quantified

We directly compared expression levels corresponding to conditions of Figures 2F and 2H using a monoclonal human a-syn antibody. We detected no a-syn labeling above background in pcDNA transfected cells (Figure S4A, B), whereas Wt a-syn labeling is clearly visible for conditions corresponding to low expression levels (Figure S4C), and substantially brighter for those corresponding to high expression levels (Figure S4D). These images also demonstrate that >90% of all cells expressing VAMP8-pHluorin also express a-syn (Figure S4C, D). We quantified differences in expression levels of Wt, A30P, A53T, and G51D with flow cytometry (as gated by VAMP8-pHluorin expressing cells), which showed a mean fluorescence 3-4-fold higher for cells transfected with the higher concentration of construct DNA compared to those transfected with the lower concentration (Figure S4 E, F).

We further employed western blots to estimate the absolute amounts of Wt a-syn expressed in RBL cells after transfection with the lower and higher amounts of DNA, which we calibrated using purified samples of recombinant a-syn (Figure S5). Converting to cellular concentrations, we determined the low expression level to be 130 μg/ml or 9 μM and the high expression level to be 300 μg/ml or 20 μM. These values fall within the range reported for concentrations of a-syn found in living neurons (5-50 μM or 75-750 μg/ml).^40^

### Low expression of human a-syn inhibits stimulated exocytosis in PC-12 cells

We tested the generality of our results using PC12 cells, which are derived from the adrenal medulla, secrete dopamine, and have been used as a model system for neurosecretion. Transfected cells expressing VAMP8-pHluorin and pcDNA show 25% exocytosis stimulated by thapsigargin (Figure S2A). This stimulated response is inhibited by 70% for cells transfected with Wt a-syn expressed at low concentrations (Figures S2B, C), consistent with our results with RBL cells (Figure 2F).

### Stimulated endocytosis is inhibited at higher expression levels of Wt a-syn, depending on its C-terminal tail

Several studies implicating a-syn as an important regulator of synaptic vesicle trafficking have focused on the endocytic events,^41–44^ and we evaluated effects of a-syn on stimulated endocytosis in RBL cells. Cells were transfected with mRFP as a reporter for positively transfected cells, together with either pcDNA or DNA for a-syn constructs at high or low concentrations. Cells were then labeled with IgE conjugated to fluorescein isothiocyanate (FITC-IgE), which displays bright fluorescence at the plasma membrane upon binding to the FceRI receptor. Crosslinking of FITC-IgE/FceRI stimulates their endocytosis, and FITC fluorescence is quenched as it moves from the neutral pH environment of the extracellular space to the more acidic environment of the endosomal compartments.^45^ Cells transfected with pcDNA and stimulated with an anti-IgE antibody exhibit fluorescence quenching due to endocytosis as monitored by flow cytometry, and the a-syn constructs we tested in direct comparison show some differences (Figure 5). We found that Wt a-syn expressed at low levels causes little or no change in stimulated FITC-IgE endocytosis (Figure 5A, E). In contrast, Wt asyn expressed at high levels causes significant inhibition (Figure 5B, E). Unlike the case with stimulated exocytosis (Figure 2H) A30P expressed at high levels inhibits stimulated endocytosis similarly to Wt a-syn (Figure 5C, E). Interestingly, the truncated 1-102 mutant does not inhibit stimulated endocytosis (Figure 5D, E), suggesting that a functional C-terminus is required for this activity. Consistently, we found that Wt a-syn-mRFP, which is C-terminally tagged, also fails to inhibit stimulated endocytosis (data not shown).

**Figure 5.**
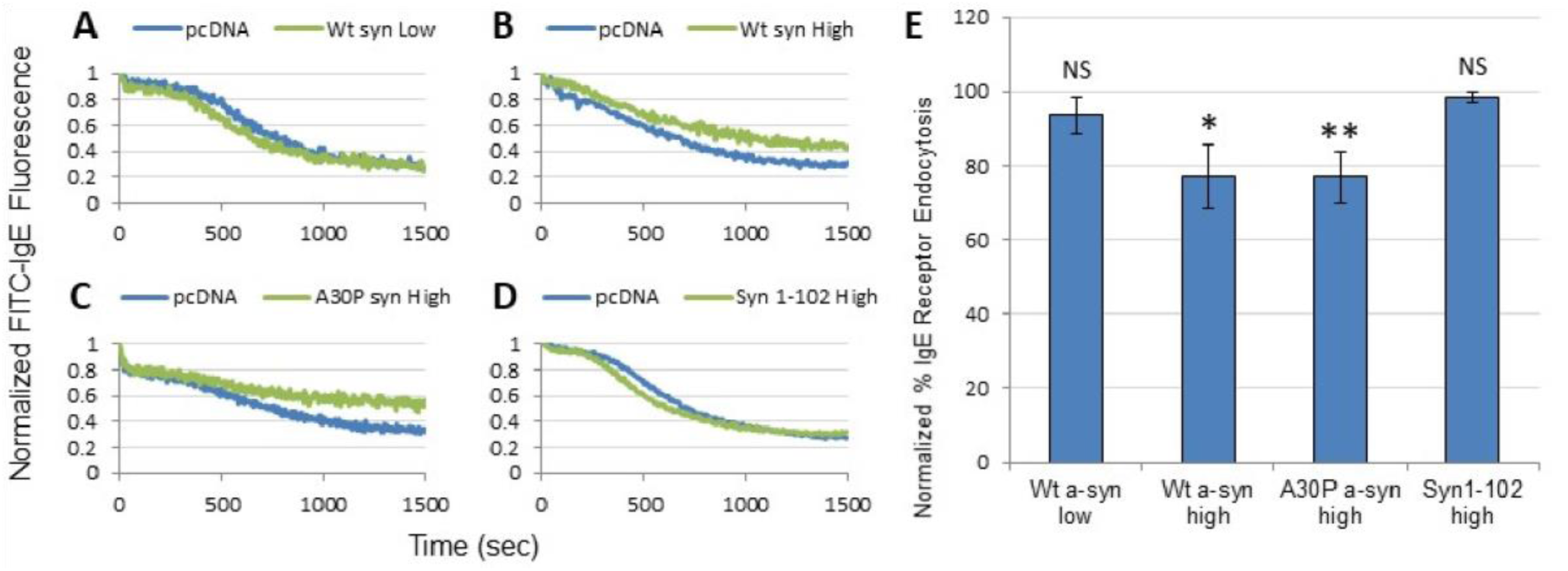
Higher expression levels of Wt a-syn inhibit stimulated endocytosis, depending on the C-terminal tail. RBL cells were co-transfected with mRFP, to mark positively transfected cells, and low levels of pcDNA or Wt a-syn (**A**), or high levels of pcDNA and Wt a-syn (**B**), A30P a-syn (**C**), or 1-102 a-syn (**D**). Cells were sensitized with FITC-IgE, incubated at 37°, and stimulated with anti-IgE antibody. Endocytosis of FITC-IgE was monitored in a flow cytometer by FITC fluorescence quenching. Fluorescence values prior to stimulation were set to 1.0, and fractional fluorescence after stimulation were compared directly to the pcDNA sample run in parallel, corresponding to maximal endocytosis. **E**) Endocytosis by indicated test samples were normalized to their paired pcDNA control samples (set at 100% endocytosis). Error bars are ± SD from 3 independent experiments. ^**^ represents P-values <0.01, ^*^ represents P-values <0.05; NS, not statistically significant (P-values >0.05).

### A-Syn at high expression levels binds to internal membranes depending on formation of N-terminal helices

We observed that cells expressing higher levels of a-syn exhibit intracellular puncta that can be visualized with fluorescent immunolabeling. Because a-syn is known to aggregate within neurons, we looked, but found no evidence, for a-syn aggregates in Western blots of our transfected cells (e.g., Figure S5). Overexpressed a-syn also has been reported to co-localize with various internal organelle membranes including lipid droplets,^46^ and we found that higher expression levels increase co-localization of Wt a-syn with the lipid droplet marker Nile Red in RBL cells (Figure S6A,B). Co-localization with lipid droplets is greatly reduced for the A30P mutant, even at high expression levels (Figure S6C), suggesting that it is driven by membrane-binding of the N-terminal region of a-syn. We also observed increased co-localization of Wt a-syn with the mitochondrial marker Mito-chameleon (Mt-cam) at higher expression levels (Figures 6F, S7), as has been previously reported for HEK cells with a high degree of a-syn over-expression^47^ and with neurons under some conditions.^48^

**Figure 6.**
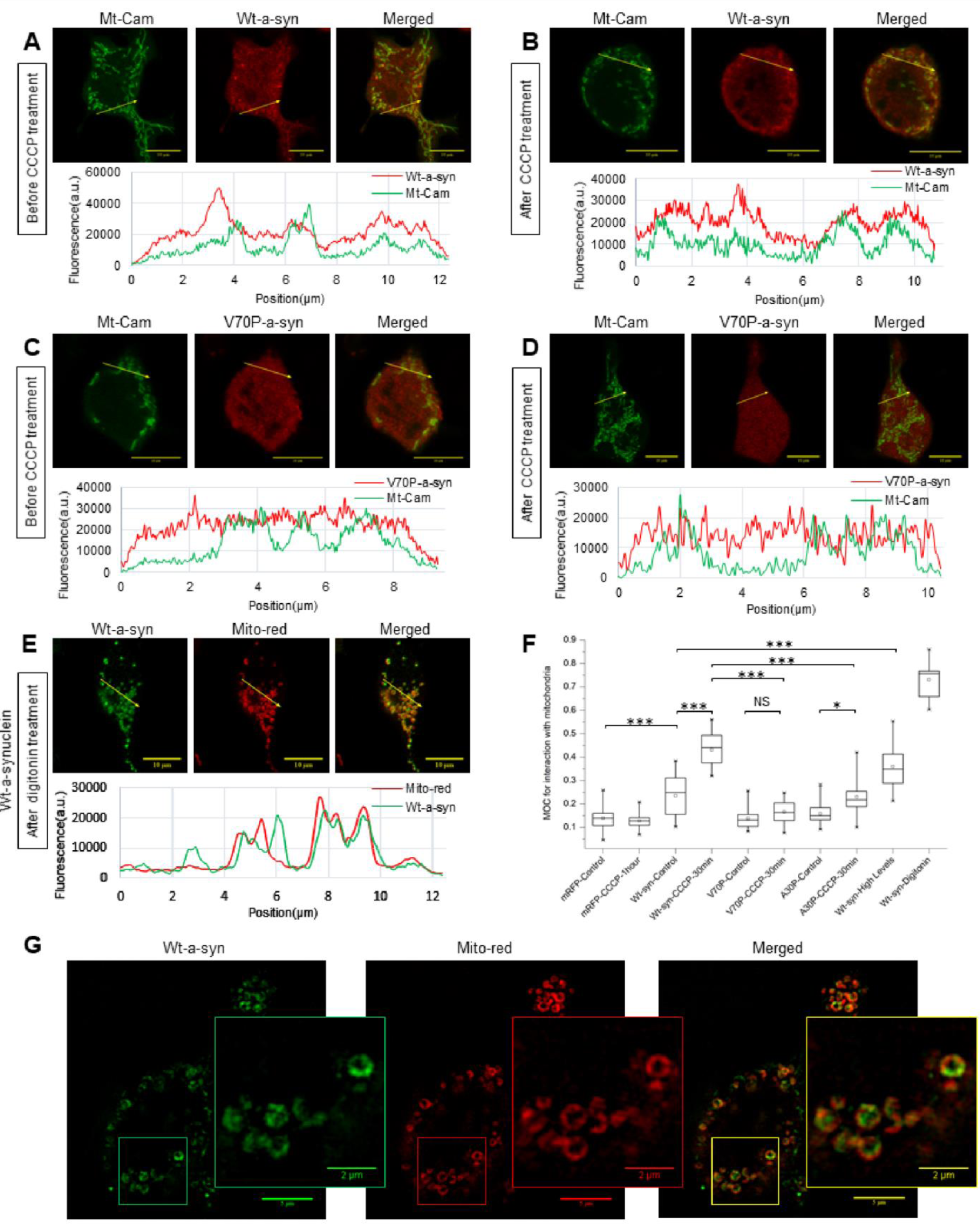
Mitochondrial and cellular stress cause association of Wt a-syn with mitochondria, disrupted by perturbation of helices 1 and 2. RBL cells co-expressing Wt asyn (**A**, **B**) or V70P a-syn (**C**, **D**) and Mito-cameleon (green) and were incubated for 30 min at 37o in BSS without (**A**, **C**) or with (**B**, **D**) 10 µM CCCP then fixed, and a-syn was immunostained (red) for visualization with confocal microscopy; scale bar = 10μm. **E)** RBL cells expressing Wt a-syn were labeled with MitoTracker Red, washed with cold PBS and permeabilized with 0.001% digitonin for 3 min. Cells were then fixed, and a-syn was immunostained (green) and visualized as above. Traces below micrographs in **A** – **E)** Fluorescence intensities across the arrow-line drawn on the images for both a-syn and mitochondrial marker channels. **F)** Averaged Mander’s overlap coefficients (MOC) for a-syn and mitochondrial marker were calculated for specified samples as represented in a box plot. The box shows 25th-75th percentile of the data; the midline shows the median, and the small square shows the average. ^***^ represents Pvalues <0.001, NS, not statistically significant (P-values >0.05). **G)** Samples prepared as described in **E** were visualized at super resolution with structure illumination microscopy, using fiduciary beads in each sample to ensure the channel alignment; scale bar is 5μm for the image and 2μm for the inset.

### A-syn co-localization with mitochondria increases after mitochondrial stress and involves helices 1 and 2

We quantified the association of transfected a-syn constructs with matrix-labeled mitochondria using both simple line scans (Figure 6A-E) and more comprehensive cross-correlation (Figure 6F). The latter approach allows a rigorous statistical analysis, and we evaluated differences using transfected mRFP as a nonspecific, baseline reference. Wt a-syn at low expression binds detectably to mitochondria, and this colocalization increases at the high expression level (Figure 6A, F). To investigate conditions that may enhance co-localization, we considered that PD-linked genes constitute the PINK1/Parkin pathway for mitophagy, which is triggered by mitochondrial stress.^49^ We tested two different conditions expected to cause mitochondrial stress: collapse of the mitochondrial membrane potential by the ionophore CCCP, and mild permeabilization of cells by low concentrations of digitonin which releases cytoplasmic ATP (as well as non-bound a-syn). We found that both of these stress conditions lead to a dramatic co-localization of a-syn with mitochondria (Figure 6B, E, F). Higher resolution, provided by structured illumination microscopy reveals that a-syn is present in regions of stressed mitochondria adjacent to, but distinct from the mitochondrial matrix (Figure 6G).^50^

We found that, compared to WT, both a-syn mutants V70P (Figure 6C, D) and A30P exhibit reduced mitochondrial localization in the absence of mitochondrial stress and a reduced increase in co-localization under conditions of CCCP stress (Figure 6F). Together with measured effects of A30P and V70P mutations on membrane interactions (Figure 3), these results indicate that intact helix-1 and helix-2 structures are involved in a-syn association with mitochondria, which increases markedly under stress conditions.

## DISCUSSION

Intracellular trafficking of membrane vesicles is observed to be abnormal in many neurodegenerative disorders, including PD,^51^ and our findings with REs in model RBL cells are consistent with a growing body of literature showing that a-syn regulates stimulated, as well as homeostatic, endocytic trafficking of transport vesicles in a variety of cells, including of SVs in neurons.^41,52–55^ Figure 7 shows simplified trafficking parallels between SVs in neurons and REs in RBL cells, based on recent reviews.^56,57^ Endosomal trafficking to sort and recycle membrane components occurs in most cell types, including neurons in which they likely intersect with specialized SV trafficking networks.^56,58–60^

**Figure 7.**
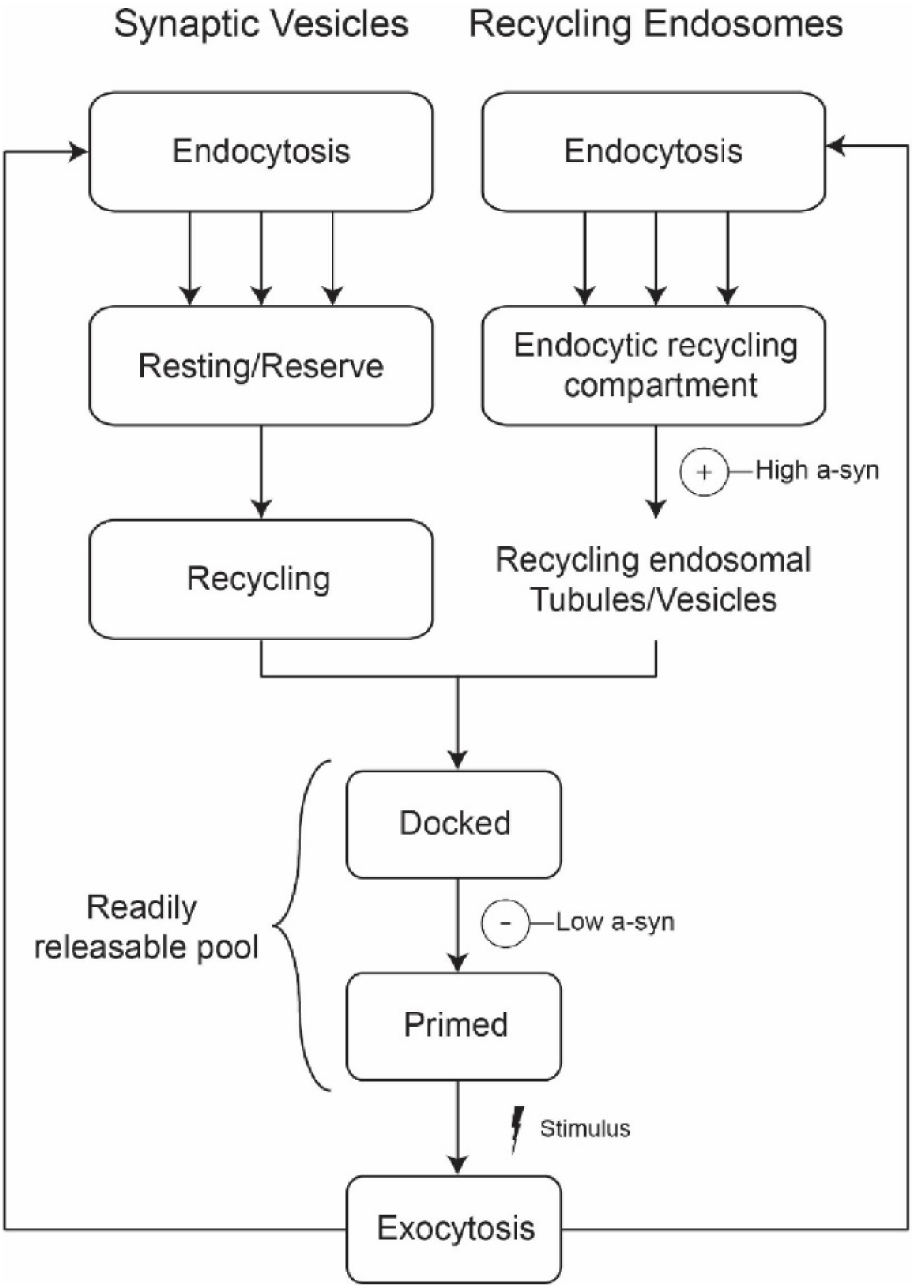
REs and SVs exhibit parallels in trafficking pathways and may be similarly affected by a-syn. Both SVs and REs are endocytosed by multiple processes and move through several endosomal vesicle stages, which we suggest have parallels. We find that Wt asyn at high expression levels increases the number of REs dispersed from the ERC toward the plasma membrane, and suggest that increased availability enhances their stimulated exocytosis (**(+)**--High a-syn); intact liposome binding is required for these effects. At low expression levels, Wt a-syn inhibits stimulated exocytosis (**(-)**--Low a-syn), depending on an intact helix-2, and we suggest the vesicles are stabilized at the docked stage, preventing priming and release; broken helix binding is required for these effects.

Both stimulated RE exocytosis in RBL cells and stimulated SV exocytosis in neurons rely upon an accessible pool of vesicles that are replenished by endocytosis. Because RBL cells contain very little (or no) endogenous a-syn (Figures S4, S5), we could monitor effects of Wt, PD-linked and other selected mutants of human a-syn, acutely transfected into these cells, to evaluate structural features of this protein as related to their *in vitro* binding properties, their binding to intracellular structures, and their functional effects (Table 1). The phenotypes we have characterized suggest that a-syn’s structural interactions with membranes as they participate in trafficking processes are similar for REs and SVs.

We observed distinctive effects depending on whether the RBL cells expressed “low” vs “high” concentrations of Wt a-syn. These concentrations, as quantified by western blotting (Figure S5), are in the range reported for neurons (5-50 μM^40,61^); comparison of fluorescent labeling intensity by flow cytometry showed our high expression level to be 3-4 times the concentration of our low expression level (Figure S4). Low expression levels of Wt a-syn inhibit exocytosis of REs, stimulated either by low-dose antigen crosslinking of IgE-receptors (Figure 2D, E), or by thapsigargin, which circumvents the receptor by directly activating downstream signaling to increase cytoplasmic Ca^2+^ (Figure 2B, F). Low expression levels of a-syn also inhibit thapsigargin-stimulated exocytosis in PC12 cells, a dopamine releasing, rat cell line (Figure S2) that has been used as an experimental proxy for neuronal exocytosis and found previously to be inhibited by a-syn.^22^

Our analysis of a panel of a-syn mutants reveals that inhibition of stimulated exocytosis at low expression levels occurs with all PD-linked mutations tested (E46K, A53T, G51D, H50Q, A30P). Inhibition also occurs with C-terminal truncation (1-102), indicating that previously described SNARE engagement or other interactions mediated by this segment are not involved in this inhibitory property.^7^ However, inhibition is eliminated by substitution of a proline residue at position 70 (V70P or A30P/V70P) (Figure 2F). This residue is located in the middle of the second of two separated helices that are formed by a-syn on the surface of detergent or lysophospholipid micelles (Figure 1C). We have postulated that this broken-helix binds simultaneously to two juxtaposed membranes, engaging docked SVs via a-syn helix-1 while binding the adjacent plasma membrane via a-syn helix-2.^17,34,36,62,63^ This double-anchor interpretation is consistent with observations that a-syn binds more tightly to membrane associated SVs than to isolated SVs.^64^ Although V70P significantly deforms the helix-2 structure in the region near the mutation site (Figure 3A,B) it has little effect on the capacity of a-syn to bind to intact isolated vesicles (Figure 3C). Therefore, we postulate that inhibition of stimulated RE exocytosis requires the interactions of intact helix-2 with the plasma membrane, which would allow specific engagement of docked vesicles via simultaneously binding of helix1. A similar double-anchor interaction of Wt a-syn has been recently proposed by others^35^ and suggested as one of several possible mechanisms contributing to inhibition of SV exocytosis in neurons by over-expressed a-syn.^14^

The A30P mutation has been shown to reduce the affinity of a-syn for SVs^65^ consistent with our measurements with synthetic vesicles (Figures 3C, S3). Because this mutation alone does not mitigate inhibition of stimulated exocytosis in our assay (Figure 2F), nor in previous reports of exocytosis inhibition in PC12 or adrenal medullary cells,^22^ we suggest that under these conditions helix 1(A30P) binding to vesicles remains strong enough to facilitate tight binding via the broken-helix bridge to the plasma membrane, as long as helix 2 is intact.

The negative regulatory phenotype we observe with Wt a-syn may represent a physiological function of a-syn in neurons. Physiologically, a-syn may participate directly in the docking process, possibly by binding first to isolated vesicles and mediating interactions with other docking factors such as Rab proteins.^27^ Notably, we found that a high antigen dose (200 ng/ml DNP-BSA) overcomes the inhibition by low expression a-syn (Figure S8) observed at the low dose of antigen (1 ng/ml DNP-BSA; Figure 2D), possibly by generating more recycling vesicles from the perinuclear ERC (Figure 7).^66^ Thus, pathological inhibition of neurotransmitter release under conditions of mis-expression may distort a finely tuned physiological role of a-syn to set an appropriate threshold for stimulation.

Our experimental system further reveals that high expression levels of Wt a-syn cause an overall increase in the level of stimulated exocytosis (Figure 2C, H). Our western blot analysis and microscopic visualization provide no evidence of a-syn aggregation that might remove an inhibitory species, pointing to a separate function of a-syn that may involve other aspects of trafficking and resulting availability of vesicles for exocytosis. We found that enhancement of stimulated RE exocytosis at higher Wt a-syn expression levels is unaffected by C-terminal truncation and all but one PD-linked mutations tested. However, the A30P mutation does abrogate this effect (Figure 2H, I), indicating that the structural requirements for this faciliatory function of a-syn differ from those of its inhibitory function. There have been previous reports of a-syn enhancement of SV exocytosis. In one study crossing CSP-alpha knockout mice, which are severely deficient in neurotransmitter release, with a-syn overexpressing mice led to a rescue, suggesting that high levels of a-syn facilitate vesicle fusion in the absence of CSP-alpha, a crucial SNARE chaperone.^67^ The A30P mutant was deficient, in agreement with our observations, but other mutants, such as a C-terminal truncation, were not examined. In another study, a-syn was found to enhance the kinetics of individual vesicle release events by altering the dynamics of fusion pore formation/closure.^68^ However, this reported effect was mitigated by the PD-linked mutations, whereas we see Wtlike enhancement by some of these mutants (A53T, G51D) in our assay (Figure 2H). To interpret our results for stimulated outward trafficking and release of REs tin RBL cells, we considered that a-syn binding to membranes is curvature dependent, with stronger binding to more highly curved membranes.^69–71^ A-syn is thought to stabilize high membrane curvature^14^ and can even tubulate membranes,^72,73^ raising the possibility that a-syn plays a role in dispersing vesicles generated via tubulation of larger membrane structures such as the ERC (Figure 7). Consistent with this, we observe a reduction in the perinuclear ERC and an increase in plasma membrane proximal vesicles in the presence of high levels of a-syn (Figure 4). We suggest that a-syn’s capacity for stabilizing membrane curvature underlies the enhancement of RE exocytosis we observe at high expression levels, changing trafficking distributions and yielding a greater availability of vesicles for docking at the plasma membrane.

To relate RE trafficking more directly to neurons we consider the SV cycle (Figure 7). As for REs, SV endocytosis replenishes pools necessary for exocytosis, and this inward trafficking process can occur through distinctive routes, some of which appear to intersect with endosomal pathways from which discrete SVs bud.^58–60^ Distinguishable SV pools have been variously described, and in Figure 7 we refer to three major pools as “Resting/Reserve,” “Recycling” and “Readily Releasable,” as described by Alabi and Tsien.^57^ Similar pools have been characterized in many other studies, in terms of their proximity to the “active zone” (AZ) of the synapse, SV clustering involving connectors or tethers, and level of stimulation required for pool activation.^14,57^ Previous studies report that expression of high levels of human a-syn prevent neurotransmitter release, in part due to inhibition of SV recycling following exocytosis, thereby reducing the recycling and readily releasable pools.^41,55,74–77^ Our characterization of asyn inhibition by broken-helix binding to docked vesicles suggests additional explanations for those previous observations, involving direct effects that impede post-docking processes such as SV priming and/or fusion.

A recent study found that SVs in the distal/reserve pool are more tightly clustered in neurons from α/β/γ-synuclein knock-out mice compared to Wt mice,^78^ with the total SV pool size remaining constant. Consistent observations were made by Nemani et al^75^ who showed in mouse brain slice neurons and in transfected neurons from the hippocampus that overexpression of a-syn disrupts SV clustering while also slightly shifting the distribution of vesicles distally from the AZ. The total SV pool was again found to be similar for over-expressed a-syn as for Wt. Their assays on neurons cultured from hippocampal and mid-brain (dopaminergic) regions showed that a-syn over-expression inhibits SV release, while a trend for enhanced release was observed when a-syn is knocked out. In this case, the functionally defined, readily releasable and recycling pools were smaller in a-syn overexpressing versus Wt neurons. This study did not explicitly describe a separate reserve/resting pool, except as the difference between the total SV pool and the other two pools. Vargas et al^78^ did not observe meaningful changes in the sizes of the readily releasable and recycling pools as defined by AZ proximity, while Nemani et al^75^ interpreted their collective results as over-expressed a-syn inhibiting stimulated SV exocytosis by limiting the access of vesicles to the recycling/releasable pool proximal to the AZ.

Despite differences relatable to distinctive cell types and vesicles, our results with REs in RBL cells are in many ways consistent with those of Vargas et al^78^ and Nemani et al^75^ and provide an expanded view. In our model, Wt a-syn promotes vesicle release from the ERC by stabilizing tubular/vesicular membranes (Figure 7), thereby increasing the pools of REs that are readily available for exocytosis at the plasma membrane. We expect that inhibitory interactions detected at low expression levels of a-syn (Figure 2) also occur at high expression levels of a-syn but that net exocytosis is enhanced because many more vesicles are available as the facilitating interactions of a-syn predominate over those that provide negative regulation. We postulate that Nemani et al^75^ observed only inhibition with over-expressed a-syn and not enhancement because synaptic terminals are very small relative to RBL cell surfaces, and SV clusters are necessarily located in close proximity to AZs. Thus a-syn driven dispersion of SV clusters appears to shift the population further from the AZ, effectively reducing the recycling and readily releasable pool. In fact, Nemani et al^75^ offered a somewhat similar explanation for their observations that the A30P mutation in a-syn abrogates inhibition of SV release in neurons but does inhibit release of chromaffin granules in adrenal medullary cells:^22^ the round shape of the latter cells (more similar to RBL cells) does not require a-syn targeting to a spatially restricted release site within axonal synaptic terminals.

Importantly, although the effect of a-syn to disperse clustered vesicles may yield different consequences for vesicle exocytosis in different cellular contexts, our results strongly indicate that this dispersive effect is a general function of a-syn that is distinct from its role in directly modulating fusion of docked vesicles, and furthermore suggest a structural basis for these different functions. Interestingly, clustering of SVs was recently proposed to be mediated by liquid-liquid phase separation of the presynaptic protein syanpsin.^79^ It is possible that the ERC, which appears as a collection of tubulated and vesiculated membranes, may also form via this recently discovered mechanism for intracellular organization. This suggests the further intriguing possibility that a-syn’s capacity to disperse SV clusters and to disperse REs from the ERC results from a-syn facilitating vesicle release from such phase-separation-induced structures.

We found that the A30P mutation reduces affinity of a-syn for vesicles and inhibits stimulated exocytosis of REs at both low and high expression levels. This reduced affinity appears to still be sufficient for tight binding via the broken-helix state to juxtaposed membranes, as we postulate above for low expression inhibition. However, this reduced affinity appears to be insufficient for stabilizing the curvature of endocytic tubules and dispersing vesicles from the ERC via the low affinity extended-helix state, which we postulate is a general mechanism by which Wt a-syn increases availability of vesicles for exocytosis. The antigen-stimulated endocytosis of IgE/FceRI we investigated in RBL cells differs somewhat from that occurring with SVs in neurons and REs in RBL cells, which both correspond to compensatory inward trafficking to recycle the exocytosed vesicles.^19^ However, key features such as curvature and fission of the invaginating membranes are likely to be similar, and a-syn may contribute to or interfere with these processes. Our observation that low expression levels of a-syn have no significant impact on stimulated endocytosis, whereas high expression levels are inhibitory and depend on the C-terminal tail (Figure 5), further demonstrates that the function of a-syn is variable with concentration and structural interactions in RBL cells. Our observations that high expression levels of Wt a-syn enhance stimulated RE exocytosis but have some capacity to inhibit endocytosis suggest the facilitating effect of increasing the pool of accessible vesicles by enhancing dispersal from the ERC is dominant.

Our search for aggregated a-syn in RBL cytoplasm at high expression levels led us to observe, instead, the association of a-syn with lipid droplets and mitochondria (Figures 6, S7), as has been reported previously.^46,47,67,80–83^ Co-localization of a-syn with mitochondria is particularly intriguing as a-syn has been reported to interfere with mitochondrial fission/fusion, transport, and autophagy in neuronal systems, and disruption of these functional dynamics is associated with PD.^48^ Indeed, the helical state of vesicle-bound a-syn is reminiscent of long amphipathic helices in mitofusions that facilitate mitochondrial membrane fusion,^84^ and a-syn’s capacities for interacting with other proteins, broken-helix binding, and curvature sensing could also affect fission/fusion and thereby other activities.

We also examined the effects of mitochondrial and cellular stress, in part because the PINK1/Parkin mitochondrial stress response pathway is strongly implicated in PD.^85^ Both treatment with the ionophore CCCP and mild permeabilization of the plasma membrane by digitonin substantially increases co-localization of low expression a-syn with mitochondria, and co-localized a-syn is distinct from the mitochondrial matrix as visualized with super-resolution imaging (Figure 6). Both a-syn mutations, A30P and V70P, cause significantly lower colocalization with and without stress treatments (Figure 6F), suggesting that membrane binding by both helix-1 and helix-2 is involved. Interestingly, a-syn has been reported to interact with the mitochondrial import machinery.^86^ During import, the inner and outer mitochondrial membranes are brought into close proximity,^87^ potentially presenting a-syn with a high affinity binding site consisting of closely apposed membranes, and requiring an intact helix-2 structure. Furthermore, the membrane composition of the mitochondrial inner membrane is rich in the negatively charged lipid cardiolipin, which may help to recruit a-syn.^88^ A-syn interactions with the mitochondrial import machinery may allow the normally cytosolic protein to cross the mitochondrial outer membrane, or a-syn may transport into mitochondria via a recently discovered mitochondrial proteostatic mechanism.^89^ Another possibility is that a-syn associates with mitochondria at mitochondrial-associated membranes, as has been observed previously,^90^ via broken-helix binding and an intact helix-2. Comparing the effects of the A30P and V70P mutations on mitochondrial co-localization to their effects on a-syn-mediated modulation of stimulated RE exocytosis suggests that a-syn first engages mitochondrial outer membranes via an extended helix (Figure 1B) and then more tightly via broken-helix binding to a juxtaposed membrane (Figure 1C).

In conclusion, we used RBL cells as a model system to study functional interactions of human a-syn, a protein with a strong link to PD but with a poorly understood function in neurological processes. These experiments allowed us to identify a regulatory role in stimulated exocytosis and endocytosis of endosomal vesicles, processes thought to be highly susceptible to neurodegeneration. We found that human a-syn can both inhibit and enhance stimulated exocytosis, depending on the expression levels of the protein. We further observed that a-syn co-localizes with mitochondria, increasing with mitochondrial and cellular stress. Intrinsic properties of a-syn binding to membranes appear to play a key role in most or all of these functions and localizations. Future experiments will aim to extend our results more directly to neurons, with the goal to clarify further the different types of membrane interactions involved in physiological function and how these are related to dysfunctional interactions that result in pathology.

## METHODS

**Cell Culture:** RBL-2H3 cells were cultured as monolayers in minimal essential medium (Invitrogen Corp, Carlsbad, CA) with 20% fetal bovine serum (Atlanta Biologicals, Atlanta, GA) and 10 *µ*g/ml gentamicin sulfate (Invitrogen) as previously described.^91^ PC-12 cells were cultured in Dulbecco’s modified eagle medium (Invitrogen) with 10% fetal bovine serum and 10 *µ*g/ml gentamicin. Adherent cells were harvested by treatment with Trypsin-EDTA (0.05%) for 8-10 min (RBL-2H3 cells) or 2-3 min (PC-12 cells), 3-5 days after passage.

**Reagents:** Thapsigargin and phorbol 12-myristate-13-acetate were purchased from Sigma-Aldrich (St. Louis, MO). Trypsin-EDTA, 0.2 µm TetraSpeck™ beads, Alexa Fluor 488-, Alexa Fluor 568-, and Alexa Fluor 647-labeled goat anti-rabbit IgG secondry antibodies were acquired from Invitrogen. Mouse monoclonal IgG^1^ anti-α-synuclein antibodies 3H2897 and 42/α-Synuclein were purchased from Santa Cruz Biotechnology (Dallas, TX) and BD Biosciences (Franklin Lakes, NJ), respectively.

**Cell Expression Plasmids:** cDNA for cell expression of human Wt a-syn, A53T a-syn, and E46K a-syn in pcDNA 3.0 vectors were obtained as a gift from Dr. Chris Rochet (Purdue). All other plasmids for cell expression of human a-syn mutants (A30P, E46K, G51D, V70P, A30P/V70P, Syn-1-102) were created within this vector by site directed mutagenesis using Phusion High-Fidelity DNA Polymerase (New England Biolabs). Plasmids for VAMP8-pHluorin and VAMP7-pHluorin were created as previously described.^23,92^ To create the Wt a-syn-mRFP plasmid, the cDNA encoding human Wt a-syn was introduced into a Clontech vector (Clontech) containing mRFP sequence, using Hind III and Kpn I restriction sites.

**Transfection by electroporation:** RBL-2H3 and PC-12 cell lines were harvested three to five days after passage, and 5 × 10^6^ cells were suspended in 0.5 ml of cold electroporation buffer (137 mM NaCl, 2.7 mM KCl, 1 mM MgCl_2_, 1 mg/ml glucose, 20 mM HEPES (pH 7.4). Co-transfections used a reporter plasmid DNA (5 µg VAMP8-pHluorin or VAMP7-pHluorin, or 1.5 µg mRFP), together with 5 µg (“low” expression) or 25 µg (“high” expression) of human Wt a-syn (or control) plasmid DNA (pcDNA 3.0, Wt, A53T, E46K, A30P, G51D, H50Q, 1-102, A30P/V70P, or V70P). For antigen-stimulated exocytosis experiments, RBL cells were cotransfected with 12.5 µg of Wt, A53T, or E46K a-syn plasmid DNA. We found higher efficiency expression for plasmids in Clontech vectors, compared to pcDNA 3.0 vectors, and correspondingly we used 1.5 µg or 10 µg of mRFP and Wt-syn-mRFP a-syn plasmids for “low” or “high” expression experiments, respectively. We find for RBL cells that cells transfected with two constructs express both or none, such that a fluorescent construct can be used as a reporter for cells co-transfected with a non-fluorescent construct.

For all conditions cells were electroporated at 280 V and 950 μF using Gene Pulser X (Bio-Rad). Then cells were immediately resuspended in 6 ml medium and cultured for 24 hr to recover; the medium was changed after live cells became adherent (1-3 hr). For exocytosis experiments the cell suspensions were added to three different MatTek dishes (2 ml/dish) (MatTek Corporation, Ashland, MA) for recovery. For antigen-stimulated exocytosis experiments, cells were sensitized with 0.5 μg/ml anti-2,4-dinitrophenyl (DNP) IgE during the recovery period.^93^

**Stimulated Exocytosis Assays:** After the electroporation recovery period and prior to imaging, cells were washed once and then incubated for 5 min at 37ºC with buffered saline solution (BSS: 135 mM NaCl, 5 mM KCl, 1 mM MgCl_2_, 1.8 mM CaCl_2_ 5.6 mM glucose, 20 mM HEPES, pH 7.4). For RBL cells VAMP8-pHluorin or VAMP7-pHluorin fluorescence was monitored for 20 sec prior to addition of either 1 ng/ml DNP-BSA, or 250nM thapsigargin, and after 6-8 min stimulation 50 mM NH_4_Cl was added. In one set of experiments, cells were first stimulated with 1 ng/ml DNP-BSA and then, after 6 min, stimulated with 200 ng/ml DNP-BSA. PC-12 cells were stimulated with 100 nM phorbol 12-myristate-13-acetate at 20 sec, and 250 nM thapsigargin at 5 min. Cells were monitored by confocal microscopy (Zeiss 710) using a heated, 40X water objective. VAMP8-pHluorin and VAMP7-pHluorin were excited using the 488-nm line of a krypton/argon laser and viewed with a 502-551 nm band-pass filter. Offline image analysis was conducted using ImageJ (National Institutes of Health). Time traces of VAMP7-pHluorin or VAMP8-pHluorin fluorescence were normalized to a 0-1 scale using the following equation: (value(t)-minimum)/(maximum-minimum), with value(t) being the measured pHluorin fluorescence at a given time, minimum being the lowest monitored fluorescent value (basal, averaged prior to stimulation), and maximum being the averaged value following NH_4_Cl addition. Percent exocytosis is calculated by the following equation: [(averaged stimulated fluorescence - basal fluorescence)/ (fluorescence after NH_4_Cl - basal fluorescence)] ^*^100.

*Total Internal Reflection Fluorescence (TIRF) Microscopy*. Samples for exocytosis were prepared as described above, and single cells were imaged in TIRF mode with a Zeiss Elyra microscope. As for the exocytosis experiments monitored by confocal microscopy, cells were stimulated with 250 nM thapsigargin before addition of 50 mM NH_4_Cl at specified time points.

**Immunostaining of a-syn variants:** Cells were electroporated and incubated in MatTek dishes for the recovery period as described for particular experiments, then fixed with 4% paraformaldehyde + 0.1% glutaraldehyde. Fixed cells were labeled in PBS with 10 mg/ml BSA using a monoclonal anti-a-syn antibody followed by an Alexa Fluor (488 or 568 or 647) conjugated secondary antibody, and then imaged by confocal microscopy or analyzed by flow cytometry.

**Distribution of recycling endosomes:** Samples were electroporated as outlined above with 5 µg of mCherry-Rab11A plasmid DNA to label recycling endosomes ^23^ and one of the following: 5 µg or 25 µg Wt a-syn, 25 µg A30P a-syn, or 3 µg or 15 µg EGFP plasmid DNA. Samples were fixed and immunostained with anti-a-syn antibody(3H2897) after the recovery period. Samples were confocally imaged using a 63x oil objective, selecting a plane near the middle of the cell including both plasma membrane and perinuclear regions. Fluorescent signal from a-syn immunostaining or EGFP was used to select the brighter cells at lower and higher expression levels, and Rab11A fluorescence was quantified to determine the relative distributions of REs using ImageJ (NIH) software. A line was drawn within ~ 800 nm of the plasma membrane, and the Rab11A fluorescence in that outer ring was divided by the total cellular fluorescence to yield the % proximal to the plasma membrane.

**Flow Cytometry and Immunofluorescence**: After transfection by electroporation, samples were plated in 60 mm dishes for standard recovery period. Cells were then harvested, washed, and resuspended in BSS. To determine expression levels, cells were fixed and labeled with a-syn antibody (42/α-Synuclein). Samples were analyzed using a BD FACSAria Fusion Fluorescence Activated Cell Sorter, and data were quantified using FCS Express 5 Flow Research software. Analysis was gated to include single cells and positively transfected cells (VAMP8-pHluorin or mRFP). For quantification of Wt or mutant a-syn expression levels Alexa Fluor fluorescence was measured on VAMP8-pHluorin gated cells. For endocytosis experiments, cells were not fixed but treated as described below; mRFP expressing cells were gated, and quenching of FITC-IgE was monitored from that subset of cells.

**Western blots to Determine a-syn Concentration in Cells:** RBL-2H3 cells were cotransfected with 5 µg of mRFP and 5µg or 25 µg of Wt a-syn plasmid DNA. After the standard recovery period cells were harvested, washed, and suspended in BSS, and sorted by flow cytometry at 37°C. ~20% of cells were identified to be transfected with Wt a-syn based on signal from the mRFP channel, and these were counted and collected for further analysis. Cells were washed in PBS, resuspended at 2×10^6^ cell equivalents/ml in lysis buffer (25 mM Tris, pH 7.4, 100 mM NaCl, 1 mM EDTA, 1% (v/v) Triton 100, 1 mM DTT, 1 mM sodium orthovanadate, 1 mM β-glycerol phosphate, 1 μg/ml leupeptin, and 1 μg/ml aprotinin), and supernatants were retained following microcentrifuge sedimentation. Standard solutions of recombinant Wt a-syn with concentrations of 2, 3.5, 5, 6.5 and 8 μg/ml were prepared by dilutions of a stock solution obtained by dissolving lyophilized, purified Wt a-syn in PBS buffer.

After filtering through a 100kDa cutoff filter, the concentration of the stock solution was determined by absorption at 280 nm using the calculated extinction coefficient of 5960 M^−1^cm^−1^. The whole-cell lysates at 7.5×10^6^ cell equivalents/ml and standard Wt a-syn solutions were resolved by SDS/PAGE (25 μl/lane), and the proteins were transferred to PVDF membranes (Immobilon-P, Millipore). After fixation with 0.4% paraformaldehyde, the membranes were blocked in 10% BSA diluted in 20 mM Tris, 135 mM NaCl, and 0.02% Tween 20 and then incubated with anti-a-syn antibody (3H2897) diluted in the same buffer. Primary antibodies were detected with HRP-conjugated secondary antibodies followed by exposure to ECL reagent (Invitrogen).

After the blots were scanned, ImageJ was used to measure the density of each band, and the amount of Wt a-syn protein in each cell lysate was interpolated from the calibration curve based on the densities of the standard solutions of purified Wt a-syn. Amount of Wt asyn per cell was calculated by dividing the amount of protein from each band by the known number of cell equivalents in the lane. Average volume of a cells was determined by transfecting RBL-2H3 cells with 5µg of EGFP and confocal imaging z-stacks (1 µm thickness) of whole cell volume. Imaris image analysis software provided a 3D rendering of cells and determined the average volume of RBL-2H3 cell to be 2627 ± 983 μm^3,^ based on measuring 50 cells.

**Mitochondrial stress assays:** *CCCP treatment*. Samples were transfected with 5μg of Wt a-syn plasmid DNA and 5μg of Mito-chameleon. After recovery cells were washed twice with BSS at 37°C, and incubated with 10µM Carbonyl cyanide m-chlorophenyl hydrazone (CCCP) for 30 min. Samples were then fixed and immunostained with a-syn antibody (3H2897) and confocally imaged using a 63x oil objective.

*Digitonin permeabilization*. Samples were transfected with 5μg of Wt a-syn, and after recovery cells washed twice with BSS buffer at 37°C, stained with 200 nM MitoTracker^®^ Red CMXRos at 37°C for 30 min, followed by three washes with BSS buffer at 37°C. Cells were then washed with cold PBS and permeabilized with 0.001% digitonin in cytosolic buffer (15 mM HEPES, 50 mM PIPES, pH=6.9, 1 mM MgSO^4^, 4 mM EGTA, 2 mM DTT, 1 μg/ml each of leupeptin, and aprotinin) for 3 min, then washed carefully with PBS. After fixing, samples were immunostained with a-syn antibody (3H2897) and confocally imaged using a 63x oil objective. For super resolution images of mitochondria within digitonin permeablized cells, structure illumination microscopy was used. Images were acquired on a Zeiss Elyra microscope utilizing a 63x oil objective, and 0.2 µM TetraSpeck™ beads were used as fiducial markers to ensure alignment of different imaging channels.

*Analysis of a-syn association with Mitochondria*. 20 cells from three experiments for each condition (with and without stress) were analyzed as follows. The confocal image was split to a-syn and mitochondria fluorescence channels, and the fraction of signal from a-syn channel which overlaps with the signal from mitochondria was quantified as the Manders overlap coefficient using ImageJ via the JACoP plugin.

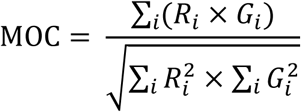

**Stimulated Endocytosis Assays:** RBL cells were co-transfected with 5 µg of mRFP and 5 µg of pcDNA or 5 µg Wt a-syn plasmid DNA (low a-syn expression level) or 5 µg of mRFP and 25 µg of pcDNA or 25 µg of Wt a-syn, A30P a-syn, or 1-102 a-syn plasmid DNA (high a-syn expression level). After the recovery period, cells were harvested, washed and suspended in BSS, and incubated with 3 µg/ml FITC-IgE for 45 min at 37°C. After washing, IgE-sensitized cells were analyzed by flow cytometry at 37°C: IgE/FcεRI complexes were crosslinked by addition of anti-IgE antibody at t = 0 sec. Acidification of internalized complexes was monitored by FITC fluorescence quenching, gated on cells positively expressing mRFP. Data were collected for ~1500 seconds, and analyzed with FCS Express 5 Flow Research software.

**Fluorescence detection of a-syn association with lipid droplets:** Samples were transfected with 5 µg of Wt a-syn, 25 µg of Wt a-syn or 25 µg of A30P a-syn plasmid DNA, and after recovery washed twice with BSS at 37°C. Cells were covered to protect from ambient light and incubated with 1ml of Nile Red solution (1ng/ml in 150 M NaCl) for 10 min at room temperature. After washing twice with 2 ml PBS and fixing, cells were immunostained with asyn antibody (3H2897). Z-stack confocal images of whole cells were collected with a 63x oil objective. 50 cells for each sample were imaged and quantified by counting the number of Nile Red positive lipid droplets labeled with Wt- or A30P a-syn over the whole cell volume.

**Statistical analyses for cell samples:** Statistical analyses were performed with Prism software (Graphpad) and Microsoft Excel. Statistical significance was determined by a OneWay ANOVA (Analysis of Variance) followed by Tukey's post hoc test using Origin software.

Level of significance is denoted as follows: ^*^*P* < 0.05, ^**^*P*< 0.01, ^***^*P* < 0.001.

**Expression and purification of recombinant isotope-labeled a-syn for NMR:** Wt human a-syn cloned into a pT7-7 vector and a-syn mutants (A30P, V70P, A30P/V70P and C-terminal truncation (1-102)) created using site-directed mutagenesis (Agilent QuikChange) were confirmed by Sanger sequencing. E. coli BL21 (DE3) cells were transformed with plasmid DNA and grown in M9 minimal media supplemented with ^15^N-ammonium chloride and ^13^Cglucose as the sole nitrogen and carbon sources, respectively. Recombinant protein production was induced by addition of IPTG at mid-log-phase growth period (OD_600_=0.6), and the cells were harvested 2-3 hours after induction.

Cell pellets of full-length variants were lysed by sonication, followed by ultracentrifugation, acid precipitation of the resulting supernatant at pH 3.5 and precipitation by addition of 50% ammonium sulfate as previously described.^94^ The purified protein was lyophilized following dialysis into water. Truncation mutants are not acid-stable, and 1-102 asyn was purified subsequent to cell lysis and ultracentrifugation using a series of ammonium sulfate cuts followed by anion exchange chromatography and reverse phase HPLC prior to lyophilization.^30,95^ Purity of the samples was assessed by SDS-PAGE, and subsequent NMR spectra provided additional confirmation of sample purity.

**NMR Sample preparation:** Lyophilized proteins were dissolved in NMR buffer (10mM Na_2_HPO_4_, 100mM NaCl, 10% D_2_O, pH 6.8) at an initial concentration of approximately 300 µM and filtered through a 100kDa cutoff filter to remove possible higher molecular weight oligomers. Protein concentration after filtration was assayed by absorbance at 280 nm using the calculated extinction coefficient of 5960 M^−1^cm^−1^ for full-length variants. In addition to absorbance measurements, the integrated intensity of the amide proton region in 1dimensional NMR ^1^H-^15^N HSQC spectra was used to generate samples of different a-syn variants at equal concentrations. Protein solutions were mixed with lipid or detergent stocks to generate vesicle- or micelle-bound samples.

**NMR of SDS micelle-bound a-syn:** Samples of full-length or C-terminally truncated asyn variants were mixed with deuterated (d25) SDS stock solution to a final SDS concentration of 40mM. Triple resonance experiments (HNCA, HNCACB, CBCACONH) were conducted at 40°C to transfer peak assignments for resonances that moved relative to the assigned Wt asyn spectrum and to determine the CA and CB chemical shifts.^96,97^ Data were collected on a 600 or 700 MHz Bruker Avance spectrometers equipped with cryogenic probes and located at the Weill Cornell NMR Core facility or at the New York Structural Biology Center. Sensitivity improved versions of the NMR experiments employing pulsed-field gradient were used.^98^ Typically for 3D experiments, 512 complex points were acquired in the ^1^H dimension centered on the water frequency (~4.7 ppm) with a spectral width of 10 ppm, 32 complex points were acquired in the ^15^N dimension centered at ~118 ppm with a spectral width of 26 ppm, and 64 complex points were acquired in the ^13^C dimension with the center frequency and spectral window optimized to increase resolution without aliasing the peaks. Spectra were processed using NMRPipe^99^ and visualization for sequential assignment was performed using CCPNmr Analysis.^96^ Spectra were referenced indirectly to DSS (2,2-dimethyl-2-silapentane-5-sulfonate) and ammonia using the temperature-adjusted chemical shift of water. For full-length V70P asyn samples, the carbon dimension in triple-resonance spectra was re-referenced using the Wt a-syn protein by minimizing the average difference between the mutant and Wt chemical shifts for residues 111 to 130.

Amide group chemical shift deviations from Wt a-syn were calculated as

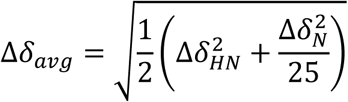

where, ∆*δ_HN_* is the amide proton chemical shift difference in ppm and ∆*δ_N_* is the amide nitrogen chemical shift difference in ppm

Secondary carbon chemical shifts were calculated as

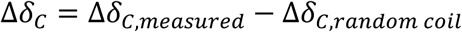

Where, ∆*δ_C,measured_* is the measured chemical shift, and ∆*δ_C,random coil_* is the sequence and temperature-corrected random coil chemical shift.^100,101^

**Lipid SUV vesicle binding assay:** Lipid SUVs were prepared by sonication as described previously^97,102^ using a lipid mixture of DOPC: DOPE: DOPS at a molar ratio of 60:25:15, dried, hydrated using NMR buffer to a lipid concentration of 20 mM, sonicated until clarification, and ultracentrifuged to remove non-SUV membranes. This stock solution was mixed with a-syn protein stock solutions at defined ratios to generate samples for NMR spectroscopy at 5 and 10 mM total lipids. Matched lipid-free a-syn solutions were generated for comparison. Final protein concentrations were 50µM for all samples in the SUV binding experiments. NMR samples were topped with argon gas, and ^1^H-^15^N HSQC spectra were acquired at 10°C on the same day samples were prepared to preclude effects of lipid oxidation. Typically the spectra were acquired with 512 and 128 complex points, with spectral windows centered at ~4.7 and ~119 ppm, and with spectral widths of 14 and 26 ppm in the ^1^H and ^15^N dimensions respectively.

Because the SUV-bound state tumbles slowly in solution, resonances from the bound protein are broadened beyond detection. Hence, any observed peaks represent protein residues not bound to the vesicle surface. The peak intensity ratio between lipid-containing and lipid-free samples is a measure of the fraction of the protein in which a given residue is unbound, and the intensity ratio profiles can be used to compare residue-per-residue SUVbinding for different a-syn variants.

## Supporting information

## AUTHOR CONTRIBUTIONS

The experiments were designed, executed, and analyzed by MR and MMW (cell) and TD (NMR). BB, DH, and DE provided supervision and participated in the design and interpretation of experiments. The manuscript was written by all authors. *The authors have no competing interests*

## ACKNOWLEDGEMENTS

We are grateful to Dr. Alice Wagenknecht Wiesner for assistance with the western blots and thank Drs. Jeremy Dittman, Jacqueline Burre and Manu Sharma (Weill Cornell Medical College) for helpful discussions. Fluorescence imaging and flow cytometry/sorting were carried out in the Cornell University Biotechnology Resource Center with funding for the Zeiss LSM 710 confocal microscope (NIH S10RR025502) and Zeiss Elyra microscope (NSF 1428922) and with assistance from Carol Bayles. NMR measurements were carried out in the New York Structural Biology Center, which is supported by grants from NYSTAR and ORIP/NIH (CO6RR015495, S10OD018509) and the Weil Cornell NMR Core Facility, supported by grant S10OD016320 from NIH. Research support came from NSF Graduate Research Fellowship, DGE-1144153 (MMW) and NIH grants R01GM117552 (BB & DH) and R37AG019391 (DE). The content of this paper is solely the responsibility of the authors and does not necessarily represent the official views of the funding agencies.

## SUPPORTING INFORMATION

**Figure S1.**
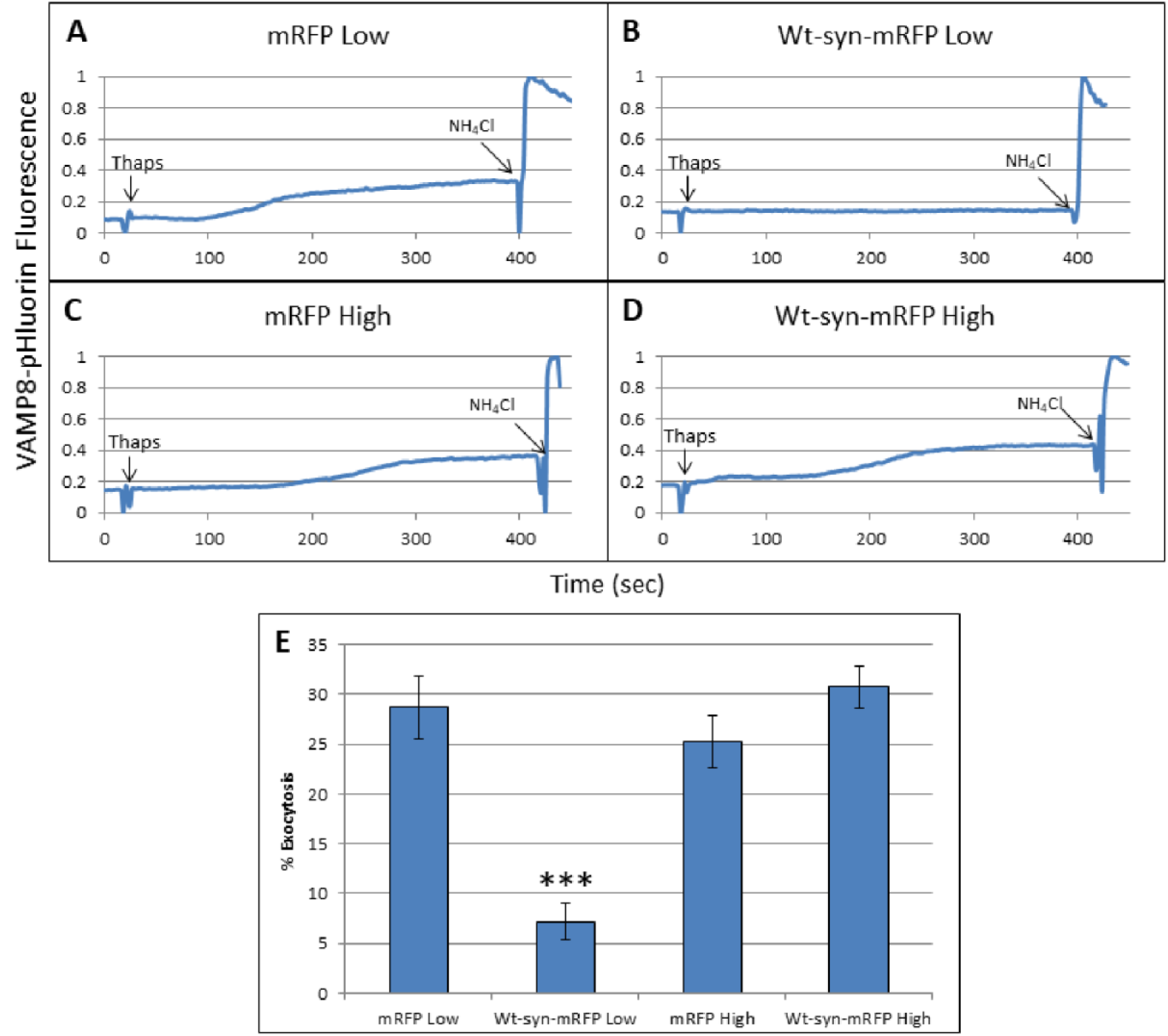
Low, but not high, expression levels of Wt a-syn-mRFP inhibit stimulated exocytosis of REs. RBL cells were co-transfected with VAMP8-pHluorin and low (**A**, **B**) or high (**C**, **D**) levels of mRFP (**A**, **C**) or Wt a-syn-mRFP (**B, D**). Exocytosis was stimulated by thapsigargin (250 nM), and after 400 sec NH_4_Cl (50mM) was added to dequench remaining intracellular VAMP8pHluorin fluorescence. **A**-**D)** Representative traces of VAMP8-pHluorin fluorescence integrated from movies of multiple confocal fields of 5-6 cells. **E)** Averaged relative exocytosis for many cells (n=55) as represented in **A**-**D**; ^***^ indicates P-values <0.001.

**Figure S2.**
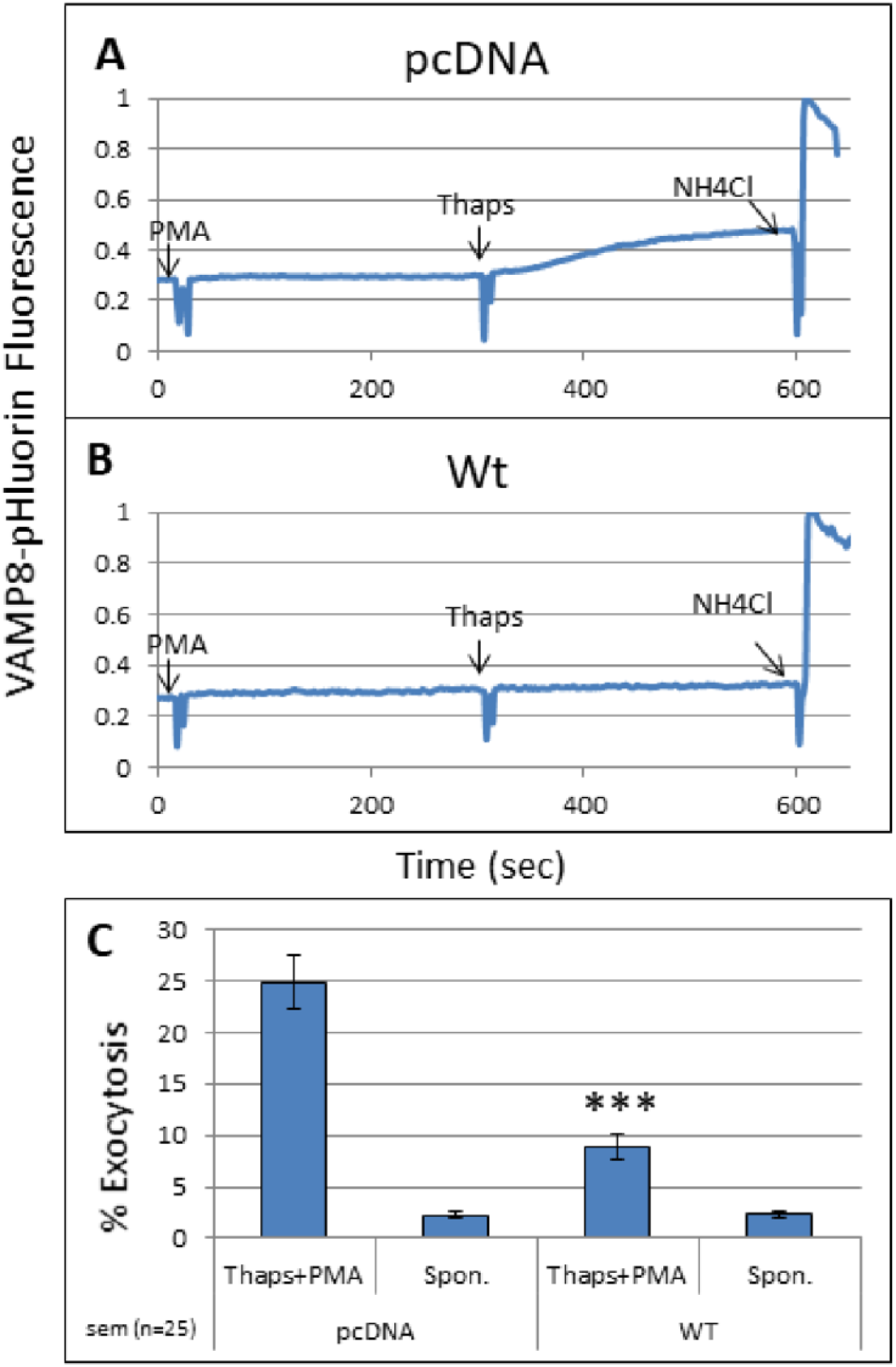
Low expression of Wt a-syn inhibits stimulated exocytosis in PC-12 cells. Traces of stimulated exocytosis averaged from 5-6 individual PC-12 cells expressing VAMP8-pHluorin and either pcDNA (**A**), or Wt a-syn (**B**). As indicated, phorbol 12-myristate-13-acetate (PMA) followed by 250 nM thapsigargin were added to stimulate exocytosis, and NH_4_Cl was added to dequench remaining intracellular VAMP8-pHluorin fluorescence. **C**) Averaged stimulated exocytosis determined from three independent experiments; error bars are ± SEM for 25 individual cells for each condition; ^***^ represents P-values <0.001.

**Figure S3.**
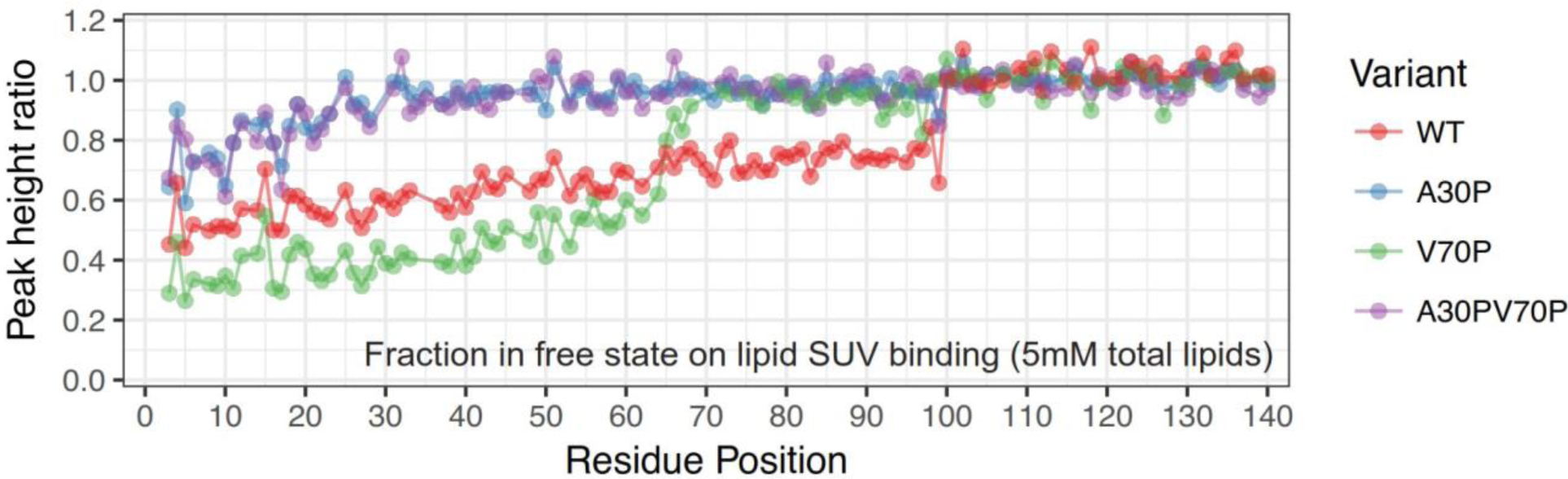
The V70P mutation results in release of a-syn regions C-terminal to the mutation site from vesicle membranes. Vesicle binding of full-length a-syn variants measured as the ratio of NMR resonance intensities in the presence and absence of liposomes. The peak intensity ratio, representing the free fraction of each residue, is plotted for 50μM protein with small unilamellar vesicles (SUVs) containing 5mM total phospholipids at a molar ratio of DOPC:DOPE:DOPS = 60:25:15.

**Figure S4.**
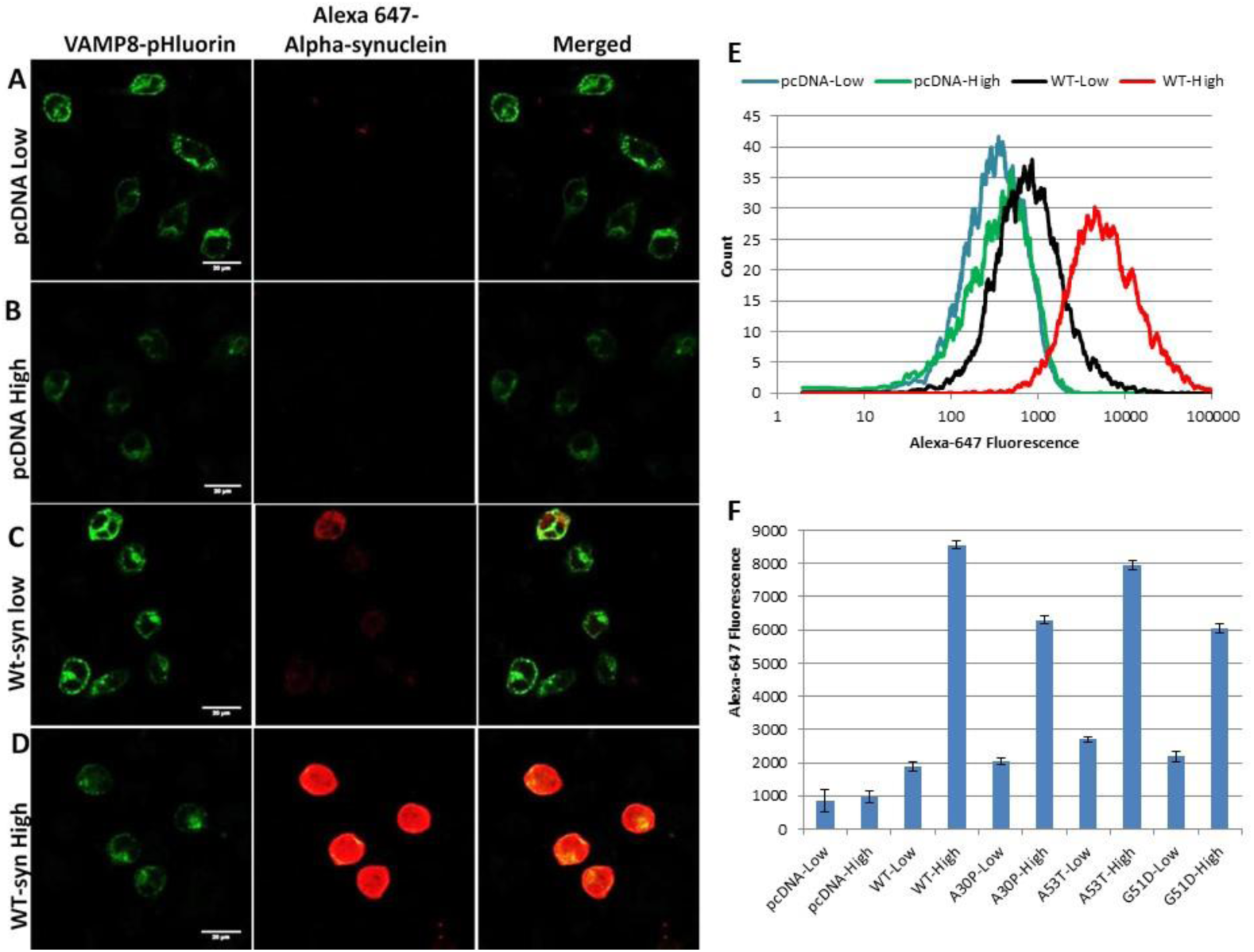
Low and high expression levels of a-syn are imaged and quantified using fluorescence microscopy and flow cytometry. **A-D)** Representative confocal images of transfected RBL cells (scale bar = 20 µm). Cells co-expressing VAMP8-pHluorin and low levels of pcDNA (**A**) or Wt a-syn (**C**), or high levels of pcDNA (**B**) or Wt a-syn (**D**) were fixed and immunostained with Alexa-647. **E**) Cells prepared as described for **A-D** were analyzed using flow cytometry in which Alexa-647 fluorescence was measured in RBL cells gated by VAMP8-pHluorin fluorescence. A representative histogram of Alexa-647 fluorescence is shown with 5,000-6,000 VAMP8-pHluorin expressing cells analyzed for each condition. **F**) Averaged flow cytometry results for low and high levels of pcDNA, Wt a-syn, and a-syn mutants, as indicated. Error bars indicate coefficient of variance for three independent experiments, with between 15,000 and 20,000 total VAMP8-pHluorin expressing cells analyzed for each sample shown.

**Figure S5.**
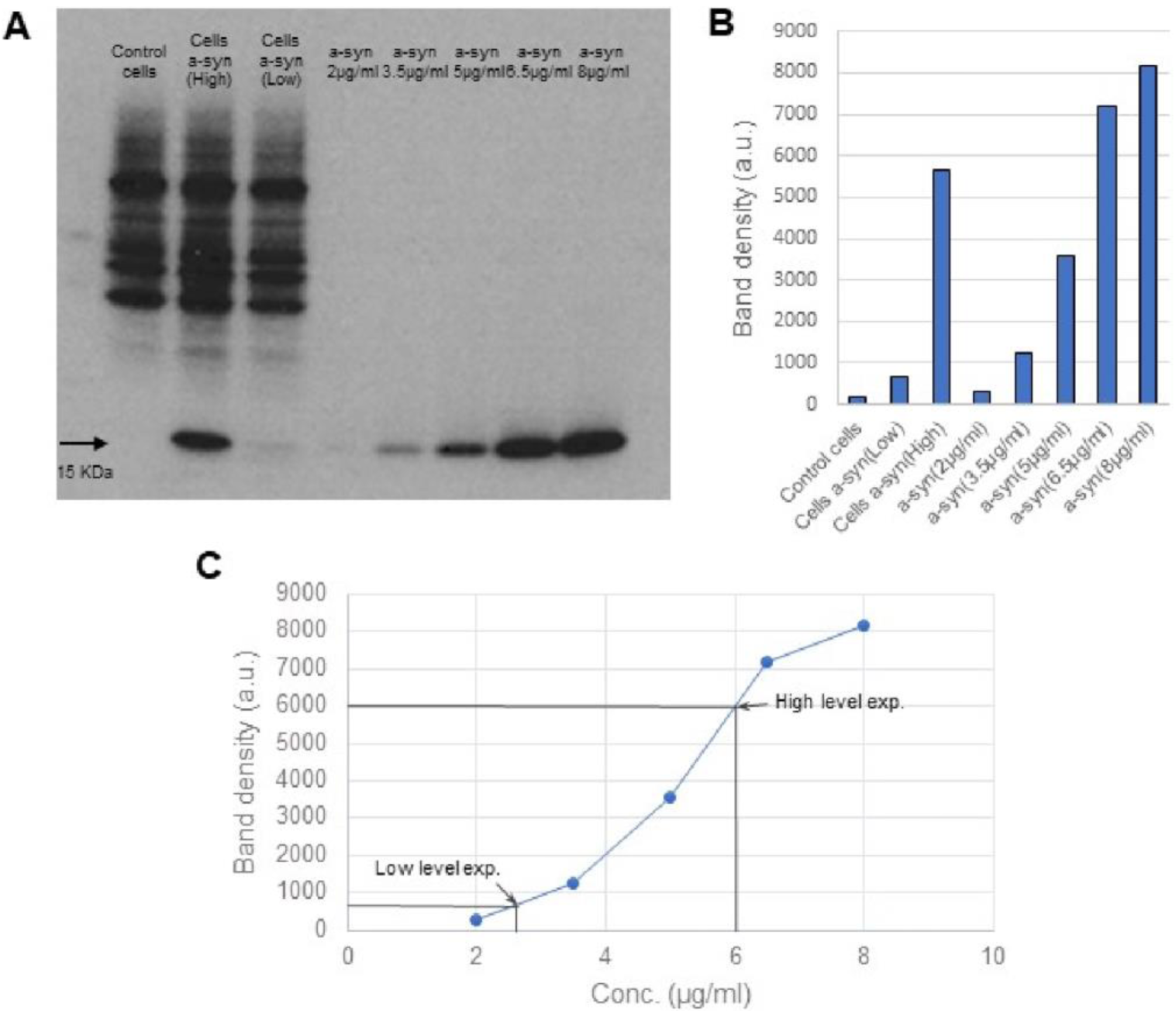
Concentration of Wt a-syn at low and high expression levels in RBL cells is estimated by western blotting. **A)** Control RBL cells and cells expressing low and high levels of Wt a-syn, co-transfected with mRFP for gating were counted and sorted by flow cytometer. Cell lysates containing 150,000 cells from each sample and standard solutions containing purified Wt a-syn at concentrations indicated were resolved by SDS/PAGE and detected by immunoblotting with anti-a-syn**. B)** The density of the band in each lane identified as a-syn was measured using the gel analysis tool in ImageJ software, and plotted. **C)** Density measurements from standard Wt a-syn solutions were used to plot the calibration curve, and the concentrations in the cell lysates were determined from this curve. The corresponding concentration in RBL cells at low (130 μg/ml) and high (300 μg/ml) expression levels of Wt a-syn was then calculated as described in Materials and Methods.

**Figure S6.**
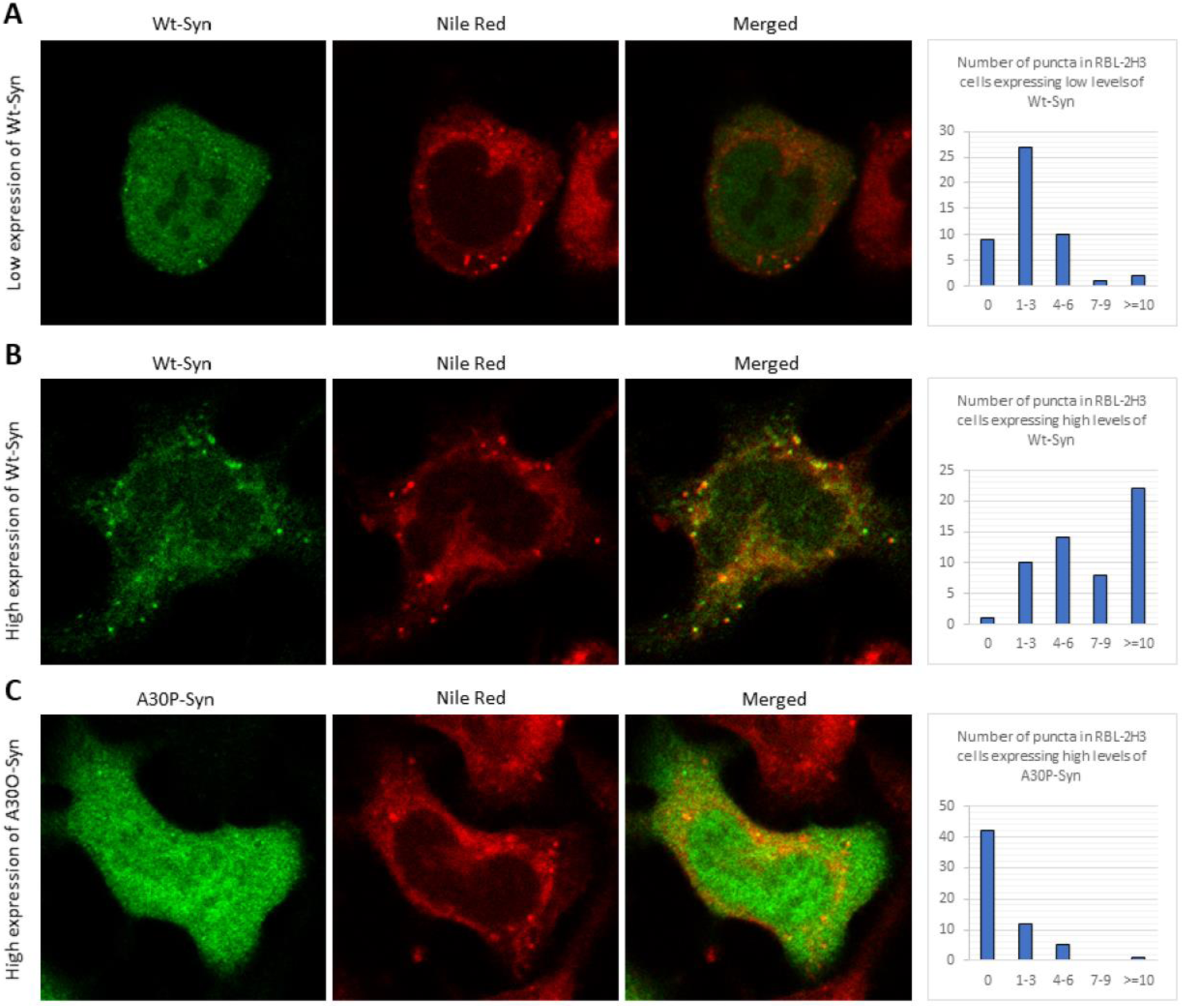
Wt but not A30P a-syn binds strongly to lipid droplets at high expression levels. Confocal images of RBL cells expressing low levels of Wt a-syn **(A)** or high levels of Wt a-syn **(B)** or A30P a-syn **(C)**. Cells were incubated with Nile red to label lipid droplets before fixing cells and immunostaining a-syn with Alexa-488. Histograms are derived from 3-D images of cells, and the number of lipid droplets decorated with a-syn was were quantified for 50 individual cells for each condition.

**Figure S7.**
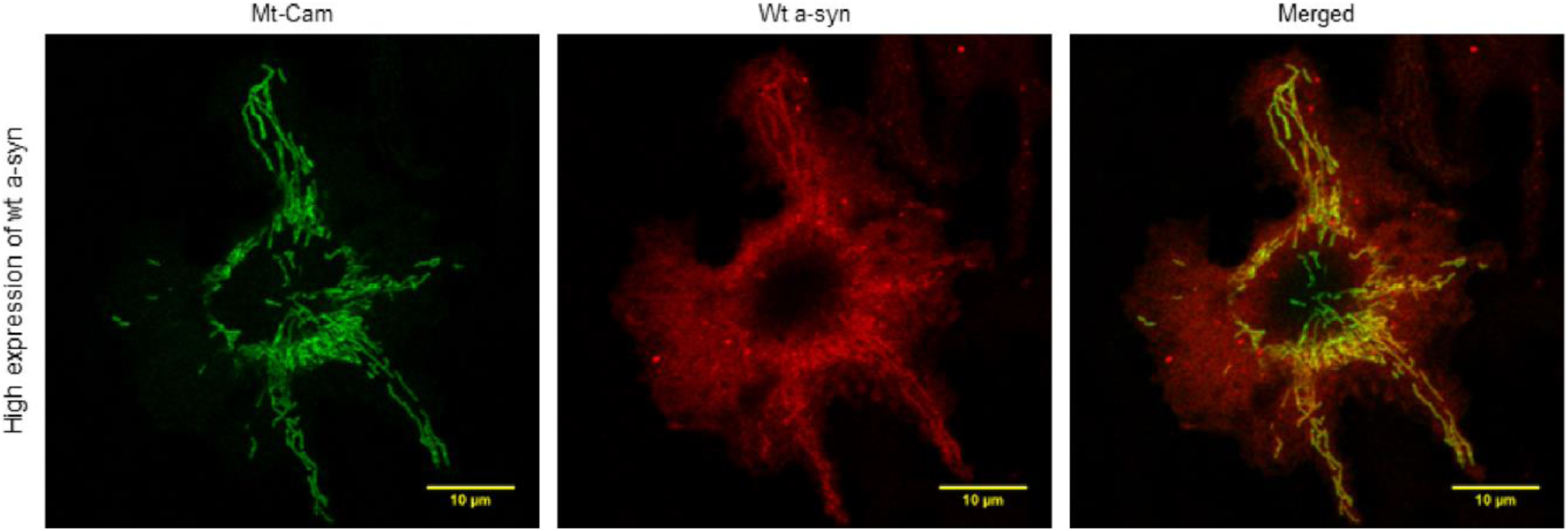
Wt a-syn at high expression levels binds significantly to mitochondria. Representative confocal images of RBL cells co-expressing Mito-cameleon and high levels of Wt asyn (immunostained with Alexa-568); scale bar = 10μm. Overlap of a-syn and mitochondrial labels for multiple cells under this condition is quantified and compared to samples with low/high levels of a-syn variants without and with mitochondrial stress in Figure 6F.

**Figure S8.**
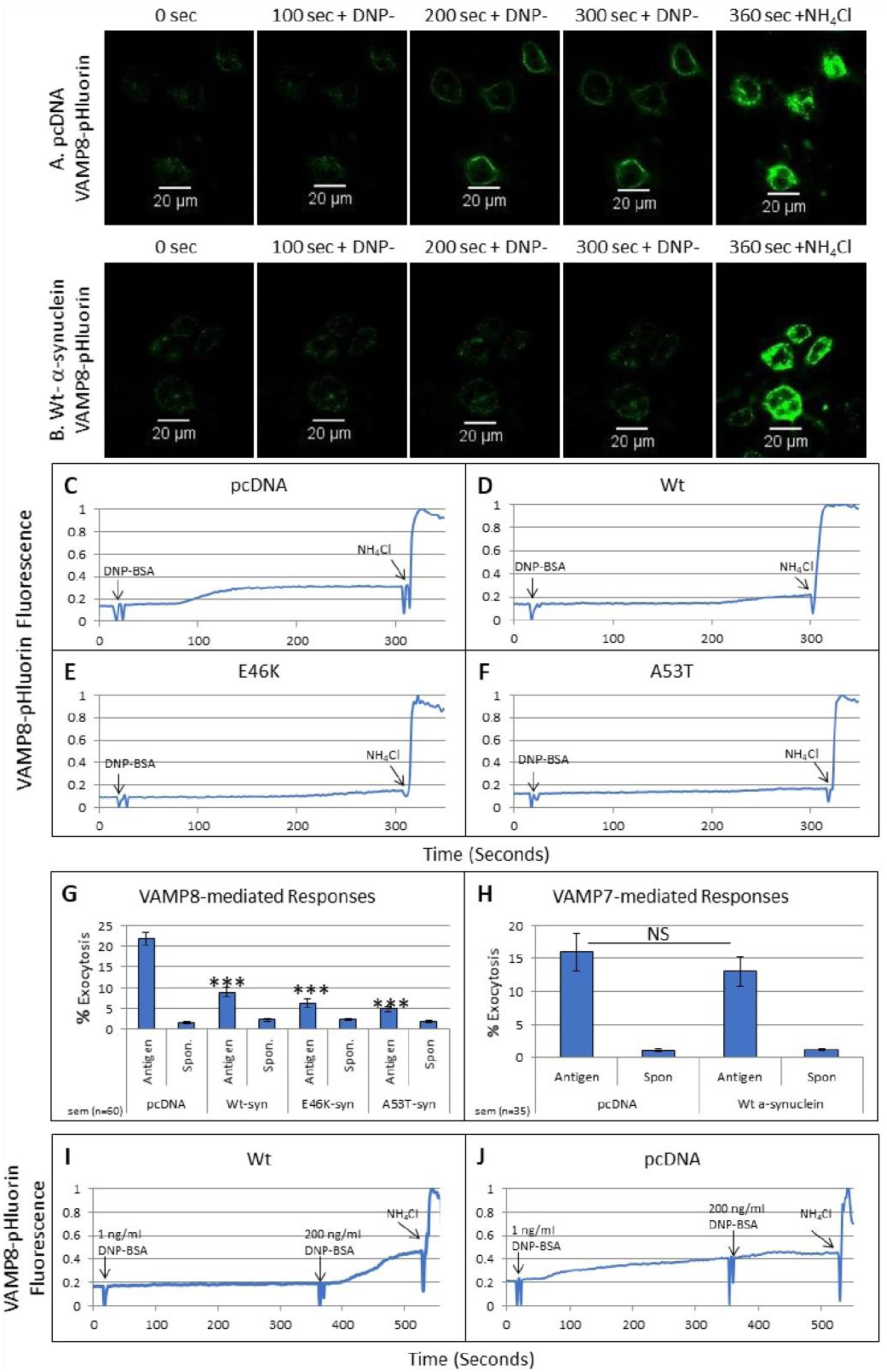
Confocal images are used to monitor exocytosis stimulated by antigen and effects of expressed a-syn variants and antigen dose. RBL cells expressing VAMP8-pHluorin and low levels of pcDNA (**A**) or Wt a-syn (**B**) were sensitized with anti-DNP IgE and stimulated with 1 ng/ml DNP-BSA at 20 sec; 50 mM NH_4_Cl was added at ~300 sec to neutralize the intracellular spaces and dequench all VAMP8-pHluorin fluorescence. Snapshots at times indicated are shown from Movies SM4 and SM5; scale bar = 20 μm **C-F**) Cells were co-transfected with VAMP8-pHluorin and pcDNA or Wt a-syn as in **A** and **B** or with low levels of E46K a-syn or A53T a-syn. Representative traces showing average change in VAMP8-pHluorin fluorescence following DNP-BSA addition are integrated from multiple fields of 5-6 cells in confocal movies, similar to Movies SM4 and SM5. **G and H**) Summary of three independent experiments monitoring fluorescence change in VAMP8pHluorin (**G**; n=40 for each sample) or VAMP7-pHluorin (**H**; n=35 for each sample) in individual cells before (spon) or plateauing after DNP-BSA stimulation, normalized to fluorescence after addition of NH_4_Cl. Error bars are ± SEM; ^***^ represents P-values <0.001, NS indicates values are not significantly different. **I-J)** RBL cells expressing VAMP8-pHluorin and Wt a-syn (**I**) or pcDNA (**J**) were sensitized with anti-DNP IgE and stimulated with 1 ng/ml DNP-BSA at 20 sec, and 200 ng/ml DNP-BSA at 360 sec. NH_4_Cl was added at 550 sec. Representative traces showing average change in VAMP8-pHluorin fluorescence were obtained as described above.

**Figure S9.**
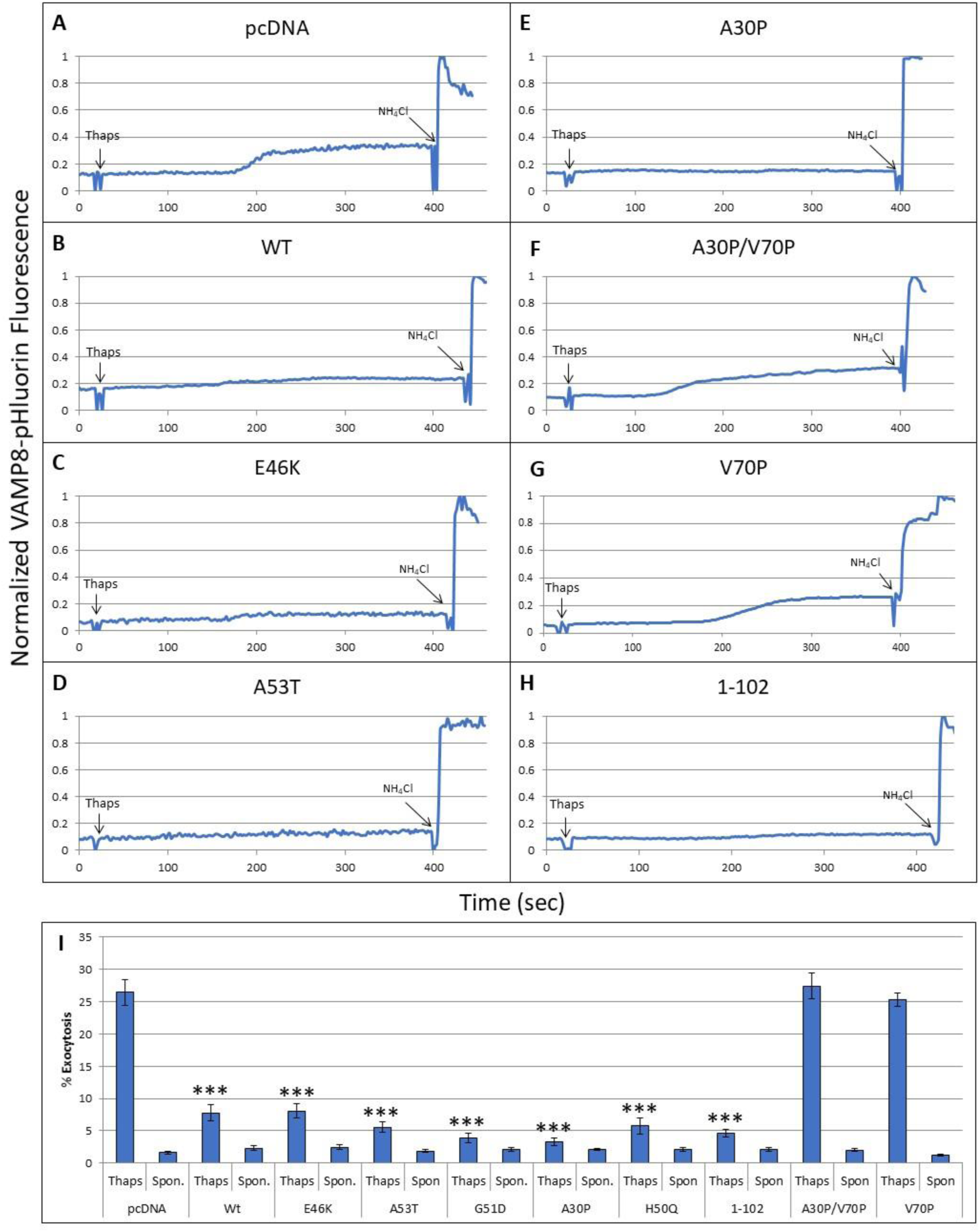
Wt a-syn and a-syn mutants except A30P/A70P and A70P expressed at low levels inhibit thapsigargin-stimulated exocytosis of REs. RBL cells were co-transfected with VAMP8pHluorin and low levels of pcDNA (**A**) or Wt (**B**), E46K (**C**), A53T (**D**), A30P (**E**), A30P/V70P (**F**), V70P (**G**) or 1-102 (**H**) a-syn. All samples were stimulated with thapsigargin at t=20 sec, followed by addition of NH_4_Cl at t= ~400 sec. Representative traces showing average change in VAMP8pHluorin fluorescence are integrated from multiple fields of 5-6 cells in confocal movies, similar to Movies SM4 and SM5. **I**) Summary of 3-4 independent experiments monitoring changes in VAMP8pHluorin fluorescence in individual cells, before (spon) or plateauing after thapsigargin stimulation, normalized to fluorescence after addition of NH_4_Cl. Error bars are ± SEM for 55 individual cells for each condition; ^***^ represents P-values <0.001.

**Figure S10.**
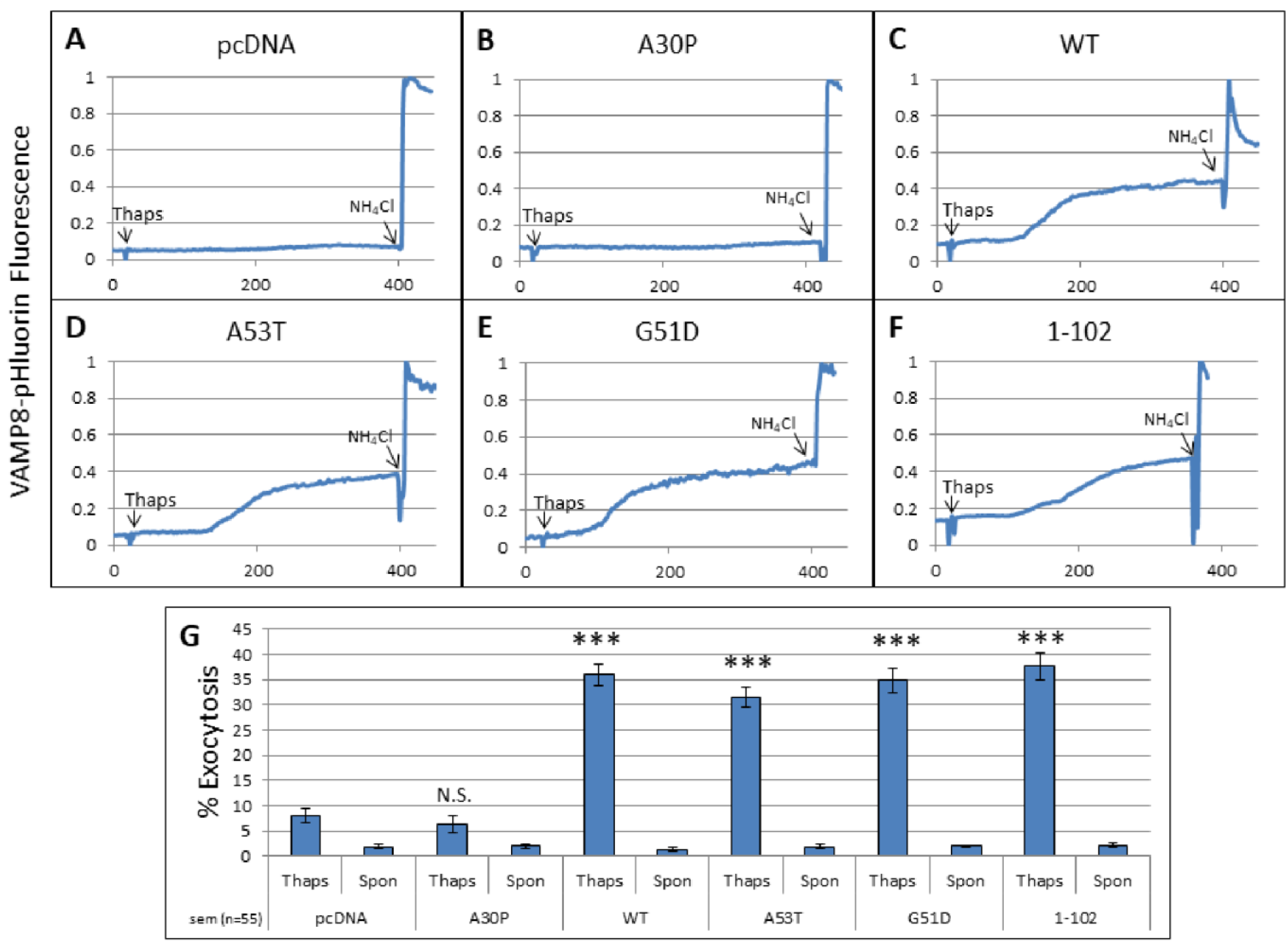
High expression of Wt a-syn and a-syn mutants except A30P enhance stimulated exocytosis. RBL cells were co-transfected with VAMP8-pHluorin and high levels of pcDNA (**A**) or A30P (**B**), Wt (**C**), A53T (**D**), G51D (**E**) or 1-102 (**F**) a-syn. Exocytosis was stimulated by addition of thapsigargin at t=20 sec, followed by addition of NH_4_Cl at ~t=400 sec. Representative traces showing average change in VAMP8-pHluorin fluorescence are integrated from multiple fields of 5-6 cells in confocal movies, similar to Movies SM4 and SM5. **G)** Summary from three independent experiments monitoring changes in VAMP8-pHluorin fluorescence in individual cells, before (spon) or plateauing after thapsigargin stimulation, normalized to fluorescence after addition of NH_4_Cl. Error bars are ± SEM for 55 individual cells for each condition; ^***^ represents P-values <0.001, N.S. not statistically significant (P-value > 0.05).

